# Genomic Islands as minimal hitchers of conjugative elements

**DOI:** 10.64898/2026.01.13.699239

**Authors:** Manuel Ares-Arroyo, Romane Junker, Amandine Nucci, Marie Touchon, Eduardo P. C. Rocha

## Abstract

Bacterial chromosomes contain Genomic Islands carrying genes involved in virulence, mutualism, and resistance to antibiotics that are transferred horizontally. Some of them conjugate autonomously (Integrative Conjugative Elements, ICEs), while others use relaxases to hitch on the former’s conjugation machinery (Integrative Mobilizable Elements, IMEs). Yet, the mobility mechanism of most Genomic Islands remains elusive. Here, we explore the hypothesis that many carry origins of transfer (*oriT*s) by conjugation. Since very few known *oriT*s were found in ICEs and IMEs, we identified 52 novel families of *oriTs* that cover most integrative elements in 6 major nosocomial species. Most of these *oriT*s are specific to integrative elements, suggesting a clear split in the targets of hitchers of conjugative elements between plasmids and integrative elements. Among 7,363 Genomic Islands, we discovered more than 1,500 IMEs carrying an *oriT* and lacking relaxases and MPF systems. These elements, coined iOriTs, form diverse large ancient families that can be found in different species. These hitchers are genetically unrelated to the putative helpers, beyond the similarity at the *oriT* sequence, and often outnumber them. They include well-known pathogenicity islands, for which the mechanisms of mobility remained unknown. Unlike plasmids, iOriTs have few antibiotic resistance genes, but high density of virulence factors. Like plasmids, the vicinity of *oriTs* concentrates defense and counter-defense systems potentially favoring Genomic Islands dissemination. Hence, iOriTs are frequent integrative mobile genetic elements that evolved to transfer horizontally by hitching on other elements while contributing to bacterial genetic adaptation.

## INTRODUCTION

Bacteria are extremely adaptable organisms that have colonized almost every ecosystem on Earth thanks to Horizontal Gene Transfer (HGT), a process in which a bacterium acquires novel traits from other genomes^1^. HGT is usually associated to self-transmissible Mobile Genetic Elements (MGEs), such as a conjugative plasmid or a bacteriophage. Autonomous conjugative elements rely on three components to transfer single-stranded DNA between contiguous cells: the relaxase (MOB), the origin of transfer (*oriT*) and the mating pair formation system (MPF). Conjugation starts when the relaxase recognizes an *oriT*, the only non-coding sequence needed *in cis* to conjugate, where it cleaves one of the DNA strands, binds to it, and carries it to the MPF. The relaxase-ssDNA is transported by the MPF into the recipient cell, where the ssDNA is circularized, and the second strand is synthesized (reviewed in ^2,3^). It was observed long ago that most plasmids are not autonomously conjugative^4^. These hitcher plasmids rely on helpers – conjugative elements – to mobilize between cells. Some encode a relaxase and an *oriT* (pMOB) and use the MPF of conjugative elements. Others exploit all the conjugation-related proteins of the latter to conjugate in a process coined *oriT* mimicry^5^. Interestingly, while most of these pOriT have an *oriT* similar to those of conjugative plasmids (pCONJ), and are thus expected to be mobilized by them, many others seem to require a relaxase from a pMOB and a MPF from a conjugative plasmid^6^. Hence, within cells there are complex networks of interdependencies between mobile elements moving by conjugation^7^. Despite receiving less attention than their autonomous counterparts, hitcher MGEs represent a common source of innovation, as they usually encode critical traits for both their bacterial host and helper element^8^.

Bacterial genomes also carry Integrative Conjugative Elements (ICE). These elements require two additional steps for conjugation: an initial chromosomal excision in the donor cell to establish a transient plasmid-like state from where conjugation starts and a final chromosomal integration step in both donor and recipient cells (reviewed in ^9,10^). For these steps, most elements rely on site-specific recombination by Tyrosine recombinases, although ICEs relying on transposable elements and or Serine recombinases have also been characterized^11,12^. The core mechanism of ICE conjugation is very similar to that of conjugative plasmids, and it was shown that most MPF types can be found both in plasmids and integrated in chromosomes^13^. While plasmid conjugation has been the focus of most past studies, the number of ICEs in bacterial genomes often exceeds those of conjugative plasmids^14^. Hitchers of conjugative elements can also integrate the chromosome and are called Integrative Mobilizable Elements (IME). Many encode a relaxase and an *oriT* similar to mobilizable plasmids^15^, henceforth called iMOB to highlight the analogy to pMOB (Fig 1A), and they were shown to be roughly as abundant as ICEs^14,16^. These elements can only transfer in the presence of conjugative elements^15^. Other IMEs encode no protein-coding genes related to conjugation, but carry an *oriT* that is similar to that of helper elements (named iOriT in analogy to pOriT). While plasmids transferring via *oriT* mimicry are abundant in model species^6,17^, few examples of iMOB and even fewer of iOriTs have been described (but see ^18–20^). As a result, the prevalence and evolutionary significance of iOriTs remains unexplored.

**Figure 1.**
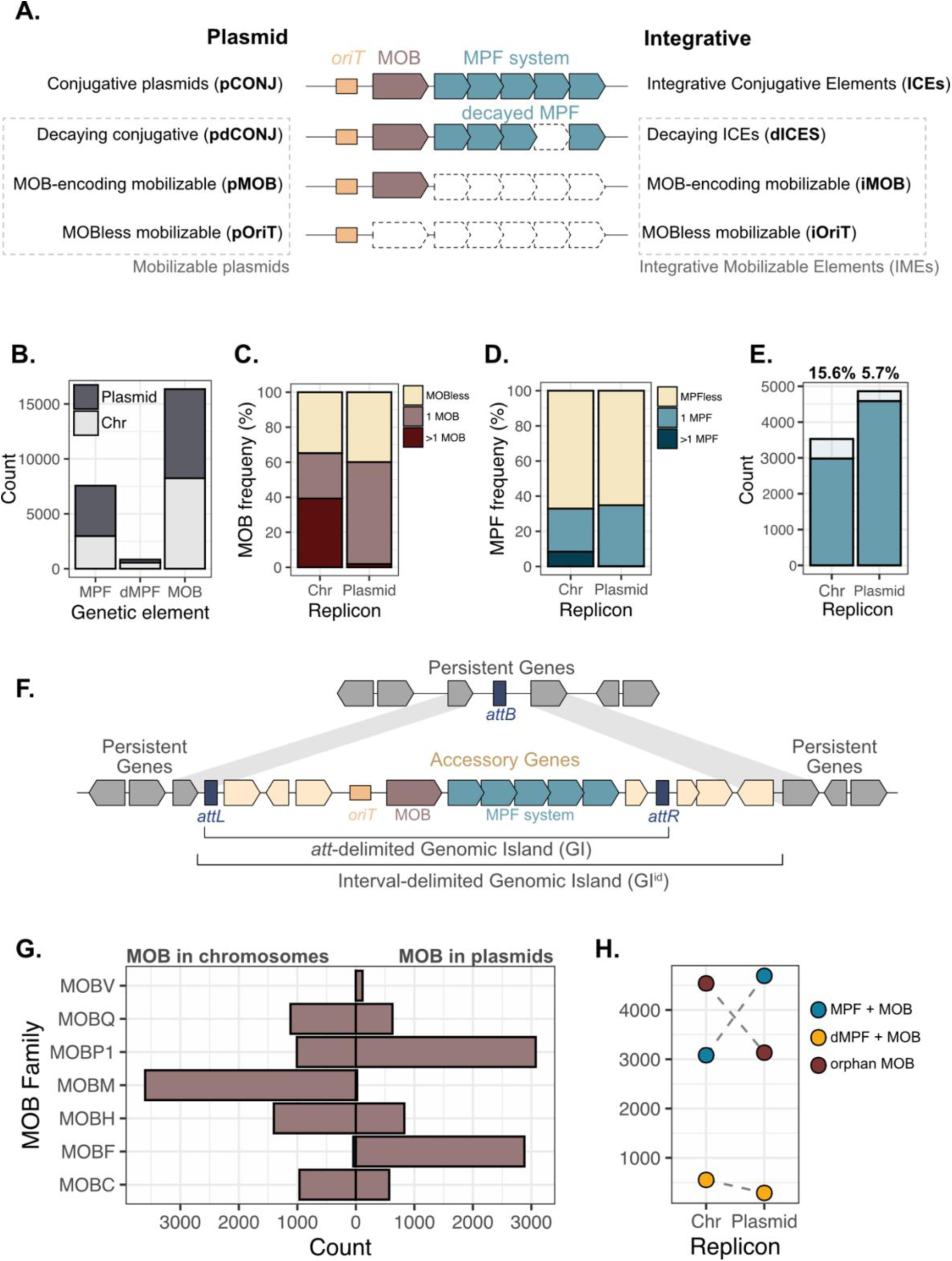
**A.** Definition of MGEs. In the middle, the genetic elements responsible for conjugation. *oriT*: origin of transfer, MOB: relaxase; MPF: mating pair formation system. At the left, MGE definition when the genetic elements are found in episomal replicons. At the right, when integrated into the chromosome. **B.** Number of genetic elements identified. dMPF: decaying MPF. **C.** Frequency of replicons carrying none, one, or more than one relaxase. Chr: chromosome. **D.** Frequency of replicons carrying none, one, or more than one MPF system. **E.** Number of complete MPF systems (dark blue) and incomplete (light blue). The number indicates the percentage of incomplete MPF systems in chromosomes and plasmids, respectively. **F.** Schematic representation of an interval-delimited and *att*-delimited Genomic Island within a spot. **G.** MOB families identified in plasmids (right) and chromosomes (left). **H.** Count of relaxases found in chromosomes (left) and plasmids (right) associated with a MPF, decaying MPF or none MPF. Grey dashed lines connect the same element in the different replicons.

Integrative mobile genetic elements provide many adaptive traits to bacteria. ICEs and IMEs spread antibiotic resistance^21^, mutualistic traits^22^, anti-phage systems^16,23^, virulence factors^24^, among many other functions. Some ICEs express these functions when dormant in chromosomes, whereas others require previous induction^25^, which can be costly^26,27^. Both IMEs and ICEs have commonly been included in the broader term of Genomic Islands (GIs), i.e., chromosomal arrays of accessory genes presumably acquired via HGT and integrated by site-specific recombination^28,29^. GIs have played a pivotal role in the emergence of critical pathogens^30^, e.g., *Salmonella* contains multiple Pathogenicity Islands^31^ among which some are conjugative^24^ or mobilizable^32^. While GIs are extremely abundant^33^, and contain may anti-phage defense systems^34^, their mechanisms of horizontal transfer are often unknown.

In this work, we wished to test the hypothesis that elements mobilizable by conjugation account for a significant fraction of the Genomic Islands in microbial genomes. We focused on six clinically relevant species of Pseudomonodota for which many genomes and MGE-related information is available. Yet, we found very few known *oriT*s in their chromosomes, which led us to identify novel families of *oriT*s by expanding a recently published comparative genomics method for the *ab initio* identification of *oriTs*^35^. We then assessed their presence across Genomic Islands to identify potential functional links between them. Finally, we devised a novel method to delimit them and could thus study the functions of their accessory genes. Our results show that iOriTs are a major class of integrative MGEs, driving the evolution of virulence and MGE-defense, and spreading horizontally thanks to a complex network of interactions with integrative and extrachromosomal elements.

## RESULTS

### Conjugative systems and relaxases are abundant in bacterial chromosomes

To identify integrative mobile genetic elements potentially transferring by conjugation, we selected six species with a critical relevance in terms of pathogenicity and antimicrobial resistance: *Escherichia coli*, *Klebsiella pneumoniae*, *Salmonella enterica*, *Pseudomonas aeruginosa*, *Vibrio cholerae* and *Neisseria gonorrhoeae*. We focused on Pseudomonodota because they have the largest number of available genomes and the best characterized conjugation systems^36^. Across the 6,182 genomes, we identified 16,340 relaxases and 8,391 MPF systems, 10% of which were classed as incomplete. Interestingly, 51% of the relaxases and 42% of the MPF systems were encoded in chromosomes (Fig 1B), despite the latter being twice less common than plasmids (6,284 chromosomes, 13,090 plasmids). Many chromosomes (41%) encode more than 1 relaxase or even MPF (8%), which is rare in plasmids (<2%, Fig 1C-D), suggesting they have multiple MGEs mobilizable by conjugation. *V. cholerae* is the only species in the dataset with two chromosomes and all MPFs were identified in the larger chromosome (Fig S1). The frequency of putatively inactivated conjugative systems is higher in chromosomes (15.6%) than in plasmids (5.7%) (Fig 1E). Most plasmids in these species are of MPF types F, T and I, of which only ~5% are incomplete, whereas chromosomes harbor MPF types G and T among which 10% and 26% respectively are classed as incomplete mostly due to the lack of the essential ATPase *virB4* (Fig S2). It is possible that defective elements integrated in chromosomes are more stably maintained in genomes than defective plasmids, which may be easily lost by segregation.

Our first analysis revealed that we were lacking the relaxase of the *Salmonella* Genomic Island 1^37^. When enquiring on the reasons of these results, we realized that the widely used programs to identify relaxases systematically miss one type of relaxase: MOB_M_. We made a novel protein profile, now added to CONJscan (see Methods), and this revealed an unprecedented abundance of ICE/IMEs with relaxases from the MOB_M_ family (not to be mistaken with the *mobM* gene in *Staphylococcus aureus* from the MOB_V_ family^38^). MOB_M_ relaxases were found to be the most common in the chromosomes, even if we could only find three previous descriptions in the literature (pCW3, SGI1 and pIGRK)^37,39,40^. MOB_M_ relaxases are distantly related to tyrosine recombinases, the existing protein profiles missed them systematically, and they were almost exclusively found in chromosomes (Fig 1G). This may explain why they were previously overlooked and occasionally annotated as integrases. The use of an accurate protein profile revealed that 96% of the MOB_M_ relaxases are located nearby an integrase (see below). They are specifically associated to MPF type G (44%, ICEs) or in iMOBs (56%). Hence, even in well-studied clades some relaxases are missed, which may result in an under-estimate of iMOB elements.

To associate the genes encoding relaxases and MPFs to integrative elements, we first identified the Genomic Islands across the genomes using the consecutive persistent gene families as delimitation. This limits MGEs to specific intervals, which can be grouped across a species in spots (see Methods), to identify similar locations across genomes^41^. Genomic Islands encoding MPF and/or relaxases defined by the edges of the intervals are henceforth called interval-delimited Genomic Islands (GI^id^) (Fig 1F, Fig S3A). In our dataset, 90.5% of the relaxases and 90.7% of MPF systems are located within GI^id^s. The remaining ones are in intervals where large rearrangements took place and cannot be assigned non-ambiguously to spots. An interval may have multiple GIs, but this should be rare since most GI^id^s encode only one relaxase (92.7%) and one MPF (>99%). In 81% of the cases, the interval with an ICE/IME had at least one integrase (68% only one) (Fig S3). Most intervals lacking an intact integrase had an integrase pseudogene (81%). This suggests that the coincidence between the MGE and the interval is high. Using this delimitation, we can separate within the chromosome the relaxases in those associated with a MPF that is complete (ICE, 53%) or incomplete (dICE, 6%), from those that are alone (iMOB, 41%). Previous works had found roughly similar numbers of ICE and IME across bacteria^14,16^ matching what is observed in plasmids^42^. Yet, in our focal species we found 1.47 times more iMOBs than ICEs (Fig 1H). The very large number of MOB_M_ relaxases may explain why we found a much larger number of relaxases than previous works. These results show that hitchers encoding relaxases are more frequent among integrative conjugative MGEs than among plasmids.

### Origins of transfer in chromosomes are different from the *oriTs* known in plasmids

Plasmids encoding no relaxases or MPF while carrying *oriTs* (i.e., pOriTs, Fig 1A) represent a large fraction of plasmids^6^. In contrast, only few IMEs lacking relaxases (iOriTs) are known to carry an *oriT*^18–20^. We searched for known families of *oriTs* and found 9,647 instances in the genomes, 83% of them in the plasmids (Fig S4). Since relaxases are expected to have a cognate *oriT* in the element and there are more relaxases in chromosomes than in plasmids, these results show that we are missing most of the *oriTs* used by the integrative elements. Indeed, while we could identify an *oriT* in most of the relaxase-carrying plasmids (~70% of pCONJ, pdCONJ, pMOB), we found very few in relaxase-carrying GI^id^s: 30% in ICEs, 15% in dICEs and almost none in iMOBs (Fig 2A). Since we missed *oriTs* even in cases where one should expect to find them (ICEs and iMOBs), this means that integrative elements and plasmids use different *oriT*s and that most of the former are unknown.

**Figure 2.**
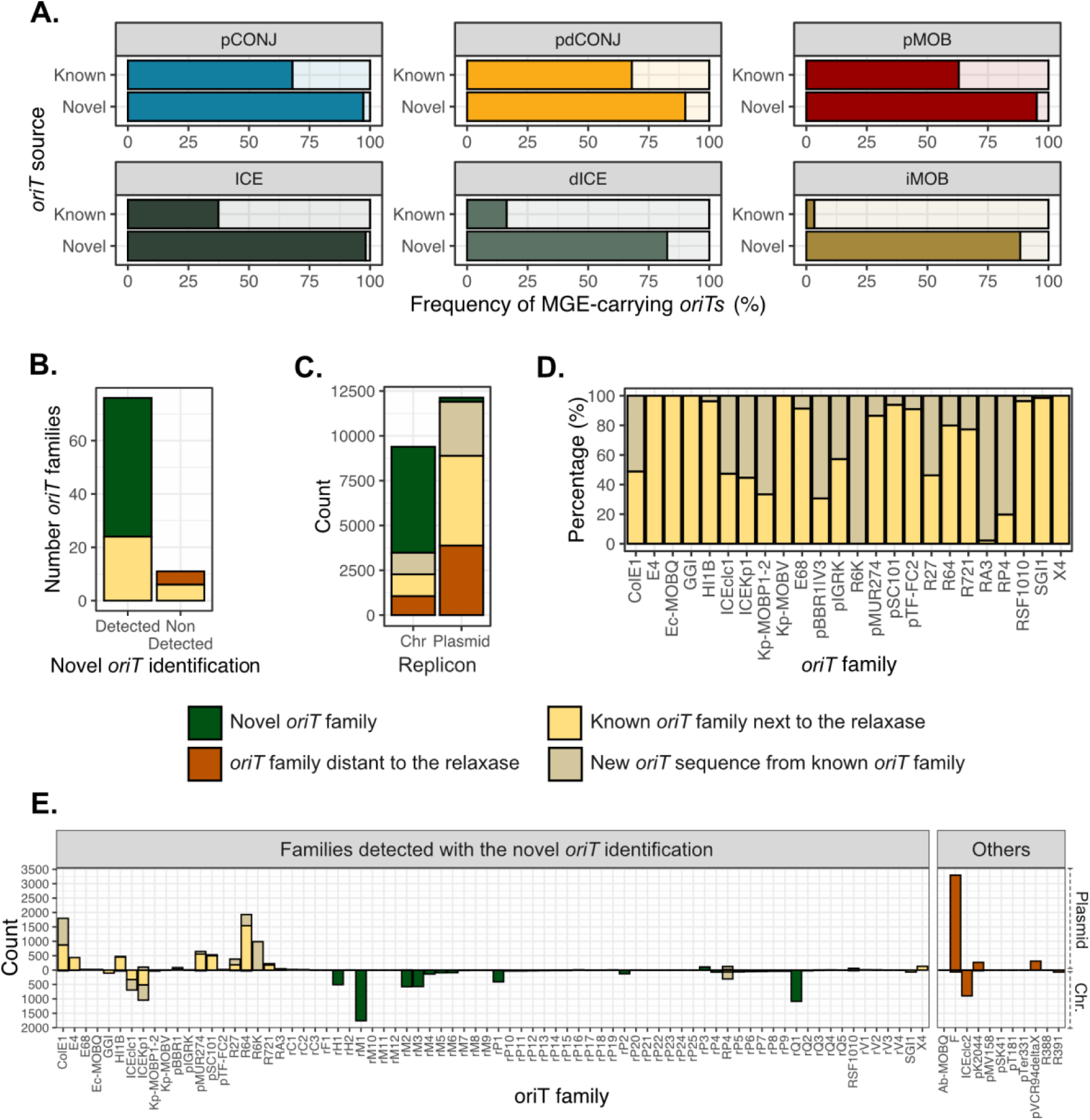
**A.** Frequency of relaxase-encoding elements with an *oriT*. Known, stands for *oriTs* already known in the literature. Novel, *oriTs* identified in this work. **B.** Number of *oriT* families detected and non-detected by our method. The color indicates the source of the *oriT.* **C.** *oriT* occurrences across chromosomes and plasmids. **D.** Known *oriT* families that were also detected by our method. The percentage indicates the fraction of new occurrences. **E.** The number of *oriT* families found in plasmids and chromosomes, the legend indicating the source of the *oriT*.

We applied our recent method to identify novel families of *oriTs* when there is no prior knowledge on their sequences^35^. Briefly, we retrieved the first large intergenic sequence upstream of every relaxase, which is the most usual location of *oriTs*, and clustered them based on their nucleotide identity while accounting for the type of relaxase (see Methods). We identified 76 candidate *oriT* families, of which a third included *oriT* sequences previously described in the literature. The remaining 52 are completely novel *oriT* families (Fig 2B), and were named according to their associated MOB family with a numeric identifier. To distinguish them from delimitated *oriT* sequences, an additional ‘*r’* standing for ‘*region’* was added to the name (e.g., *oriT*_rQ1_, *oriT*_rQ2_) (see Methods). Six previously identified *oriTs* in our first screening could not be identified by our method, presumably because they are too rare to form an *oriT* family (0.06% of the *oriTs* in our previous analysis, Fig S4). Five *oriT* families are known to be absent of the intergenic region upstream the relaxase in these species, but far from any relaxase gene, e.g. *oriT*_F_ which is next to a relaxosome-accessory protein at the opposite edge of the conjugation operon in F plasmids^43^. These *oriTs* were added manually (see Methods, Fig 2B). Finally, we built HMM DNA profiles for every known and new *oriT* candidate and used them to find occurrences in the 6,182 genomes. We identified 21,516 putative *oriTs* in both plasmids (56%) and chromosomes (44%).

We identified 4,217 *oriTs* using HMM profiles of known families, 40% of which were missed by pairwise sequence alignments (Fig 2D-E), highlighting the higher sensitivity of the HMM approach. The majority of the chromosomal *oriTs* were of the novel types (62%) (Fig 2C). The fraction of relaxase-encoding integrative elements with a candidate *oriT* increased to 93.5%, suggesting that we uncovered most *oriT*s in these species (Fig 2A). As expected, >99% of the MGEs carry the *oriT* next to the relaxase (<5Kb distance), once we accounted for the known exceptions (e.g., *oriT*_ICEclc2_, *oriT*_F_). Most plasmids (83%) and GI^id^s (85%) with an *oriT* carried just one (Fig S5), with the others often having two copies of the same family, suggesting they are in plasmid dimers or co-integrates. The similarity of trends between plasmids and GI^id^s suggests the multiplicity of *oriT*s is not caused by mis-delimitation of the latter (Fig S5A-B). Some of these cases were previously described in individual plasmids (e.g., two *oriT*_R6K_)^44^ and ICEs (e.g. *oriT*_ICEclc1_ & *oriT*_ICEclc2_)^45^. To assess if the novel *oriT*s cover most conjugative elements in a dataset independent from the one used to characterize them, we retrieved the 2,289 genomes of the six species submitted from June 2023 to May 2025. In this novel dataset, we identified 2,942 relaxases in the chromosomes and 92% of them were next to one of the *oriTs* identified (Fig S6). This suggests we covered most of the *oriT* diversity of these species.

### Integrative elements carrying only *oriTs* are frequent in bacteria

Elements carrying only an *oriT* must interact with a relaxase encoded in the helper to be transferred by conjugation^5^. Since we can now identify most MOB-associated *oriT*s, we can also identify the integrative hitchers carrying only *oriTs* (iOriT): they should have an *oriT* similar to the one of the helper carrying the relaxase. We examined the 9,930 *oriTs* located in chromosomes and found 94% within the GI^id^s (Fig S7), as expected since ~90% of relaxases and MPF systems are in these locations (see above). Around 76% were identified in GI^id^s containing relaxases (i.e., ICEs, dICEs, iMOBs) and 1,750 *oriTs* were found in GI^id^s lacking relaxases or MPF systems, i.e. in iOriT (Fig 3A). Altogether, we identified 7,363 GI^id^s corresponding to integrative elements, both ICEs, dICEs, iMOBs and iOriTs (Fig 3A). The latter often co-occur in the genomes (up to nine integrative elements in the same *P. aeruginosa* chromosome (NZ_LS998783.1)). iOriTs represent a large fraction of the IMEs (37%) and of all integrative MGEs associated with conjugation (24%). The integrative hitchers (IMEs: iOriT and iMOB) are almost double the number of autonomous elements (ICEs, Fig 3A). To note, the IME/ICE ratio varies extensively across the taxa, with IMEs outnumbering ICEs in some species (*E. coli*, *S. enterica*), while being rarer in others (*P. aeruginosa*, *N. gonorrhoeae*, Fig 3B). Mobilizable plasmids (pOriT, pMOB) are more numerous than IMEs (iOriT, iMOB) in our dataset (Fig S8). Yet, this is only true for half of the species (*Ec*, *Kp* and *Ng*), the opposite being observed in the others (*Se*, *Vc* and *Pa*), where iOriTs alone are more abundant than the whole sum of mobilizable plasmids (Fig 3B, Fig S8). In conclusion, iOriT are numerous and together with iMOB they often make the largest fraction of integrative elements transferable by conjugation, even if their abundance seems to vary between species.

**Figure 3.**
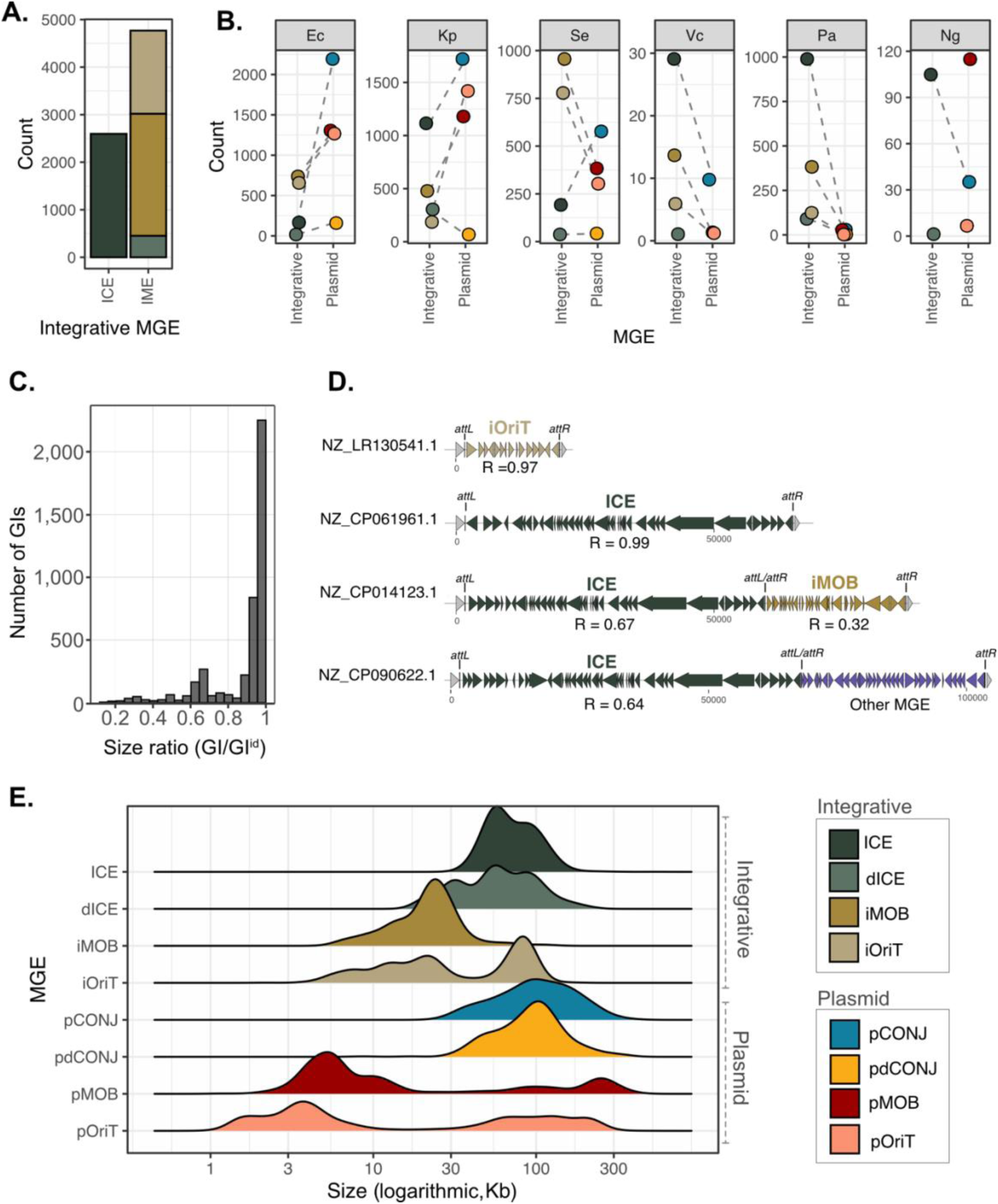
**A.** Number of integrative MGEs, both autonomous ICEs and hitcher IMEs (dIMEs + iMOBs + iOriTs). **B.** Number of integrative elements (ICE, dICE, iMOB, iOriT) compared to the number of plasmids (pCONJ, pdCONJ, pMOB, pOriT) in the six species. Dashed grey lines link MGEs carrying the same conjugative elements (ICE-pCONJ; dICE-pdCONJ; iMOB-pMOB; iOriT-pOriT). Ec: *E. coli*; Kp: *K. pneumoniae*; Se: *S. 11lasmid11*; Vc: *V. cholerae*; Pa: *P. aeruginosa*; Ng: *N. gonorrhoeae*. **C.** Size ratio of *att*-delimited Genomic Islands against interval-delimited Genomic Islands. **D.** Examples of the same genomic spot occupied by different MGEs alone or in tandem. The MGE type is annotated over the element, whereas the size ratio (GI/ GI^id^) appears under it. **D.** Size distribution of MGEs. The size is represented after a logarithmic transformation.

The log-transformed size of plasmids follows a very characteristic bimodal distribution, where conjugative elements are large and mobilizable elements are bimodal^6^. While it has been observed that the median size of conjugative plasmids is slightly higher than ICEs^46^, little is known about the distribution of size of integrative hitchers. To obtain a more precise delimitation of the elements, we searched for the elements’ *att* sites (Fig 1F). We found them for 58% of the integrative elements (4,298 out of 7,363 elements, see Methods) (Fig S9). Almost all of these *att* sites (99%) matched those recently provided in the TIGER *att* database^47^, which uses a different method to identify them, supporting the validity of our predictions (Fig S9, Table S1). In the same genomes analyzed by both our work and the TIGER database, 86% of our *att*-delimited GI match those found by TIGER even if the latter are not focusing on hitchers by conjugation (they found 95% of the ICEs in this work, but 95% of iMOBs and 98.3% iOriTs were just classed as ‘other GI’). As expected, an integrase was encoded in the 2Kb vicinity of 92% of the *att*-delimited GIs (hereinafter GIs), which in most cases encoded one and only one intact recombinase gene (82%) (Fig S9). Only 6% GIs lacked detectable vestiges of these genes. Integrases of the Serine recombinase family were very rare (<1%). GIs delimited by *att* sites are necessarily equal or smaller than those delimited by the intervals. In 80% cases, the GI retained more than 80% of the length of the original GI^id^ (Fig 3C). In cases where the GI was much smaller than the one delimited by intervals (low GI/GI^id^ ratio), additional MGEs were found integrated in tandem in the spot (Fig 3D). This indicates that intervals provide a reasonable approximation of the integrative MGEs, but that *att* site identification is important for their accurate delimitation.

The GIs show characteristic size distributions (Fig 3E). In agreement with previous works^48^, ICEs are large (mean of 75Kb) and their size varies with their MPF type: from the 100Kb MPF_F_ ICEs to the 56Kb MPF_T_ (Fig S10). Decaying ICEs are on average smaller (65Kb) than ICEs and the distribution of their sizes reveals a longer left tail suggesting that many were partially deleted (Fig 3E), as observed in plasmids with incomplete conjugative systems^49^. The iMOBs are small (23Kb) whereas the iOriTs show a bimodal distribution with many elements being more than 50kb long (Fig 3E). Inspection of the large iOriT revealed well known and precisely delimited genomic islands (Fig S11). The difference in size distribution between iOriT and iMOB is surprising, since both pMOB and pOriTs show bimodal distributions with both large and small elements^6^. While the size distribution of iOriT is the expected one, large iMOBs are rare in these species.

The present data allows to compare for the first time the size of integrative and plasmid elements transferrable by conjugation. Integrative elements show a narrower size distribution (interquartile range 22-79Kb) than plasmids (6.1Kb-116Kb), with iMOBs being on average larger than pMOBs (Kolmogorov–Smirnov test: D = 0.54, p < 2.2^−16^) and ICEs shorter than pCONJ (D = 0.35, p < 2.2^−16^). These differences are due to two effects. Among the small elements, some plasmids are extremely small (down to <1kb), whereas integrative elements are always larger than 4kb (one single 2Kb exception). This may be because the integration module requires the presence of integrases, excisionases and *att* sites, whereas plasmids only require an origin of replication and can therefore be much smaller. Among conjugative elements, the differences seem to be found at the right end tail of the distribution since some plasmids are extremely large (up to >1Mb), which is never the case of ICEs in our dataset.

### iOriTs depend on co-occurring ICEs and conjugative plasmids to transfer

The functional dependencies of IMEs, i.e., which helper MGEs they exploit, are mostly unknown. Some IMEs are known to transfer by hijacking the conjugative machinery of ICEs (e.g., the iOriT MGI*Vfl*Ind1)^50^, whereas others hijack conjugative plasmids (e.g., the iMOB SGI1)^51^. Inversely, some ICEs mobilize plasmids^52–54^. One can identify putative iOriT helpers by searching for *oriT* sequence similarity. In plasmids, *oriT* families are usually either in pCONJ (and pdCONJ) or pMOB, but not in both (Fig 4A). We found that the same applies to integrative MGEs: *oriT* families are usually associated to either ICEs (and dICEs) or iMOBs, but none is found in both (Fig 4A). Accordingly, the iOriTs usually have *oriT*s of the same families as those found in either ICEs or iMOBs, but not both (Fig 4B). Hence, while some plasmid and integrative hitchers carrying just an *oriT* for conjugation exploit both the relaxase and MPF of a conjugative element (two thirds of the cases) (Fig 4C-D), a third of them exploit a relaxase from another hitcher (iMOB/pMOB) and a MPF from a conjugative element (Fig 4C-D). The latter require the presence of two different compatible MGEs in the cell.

**Figure 4.**
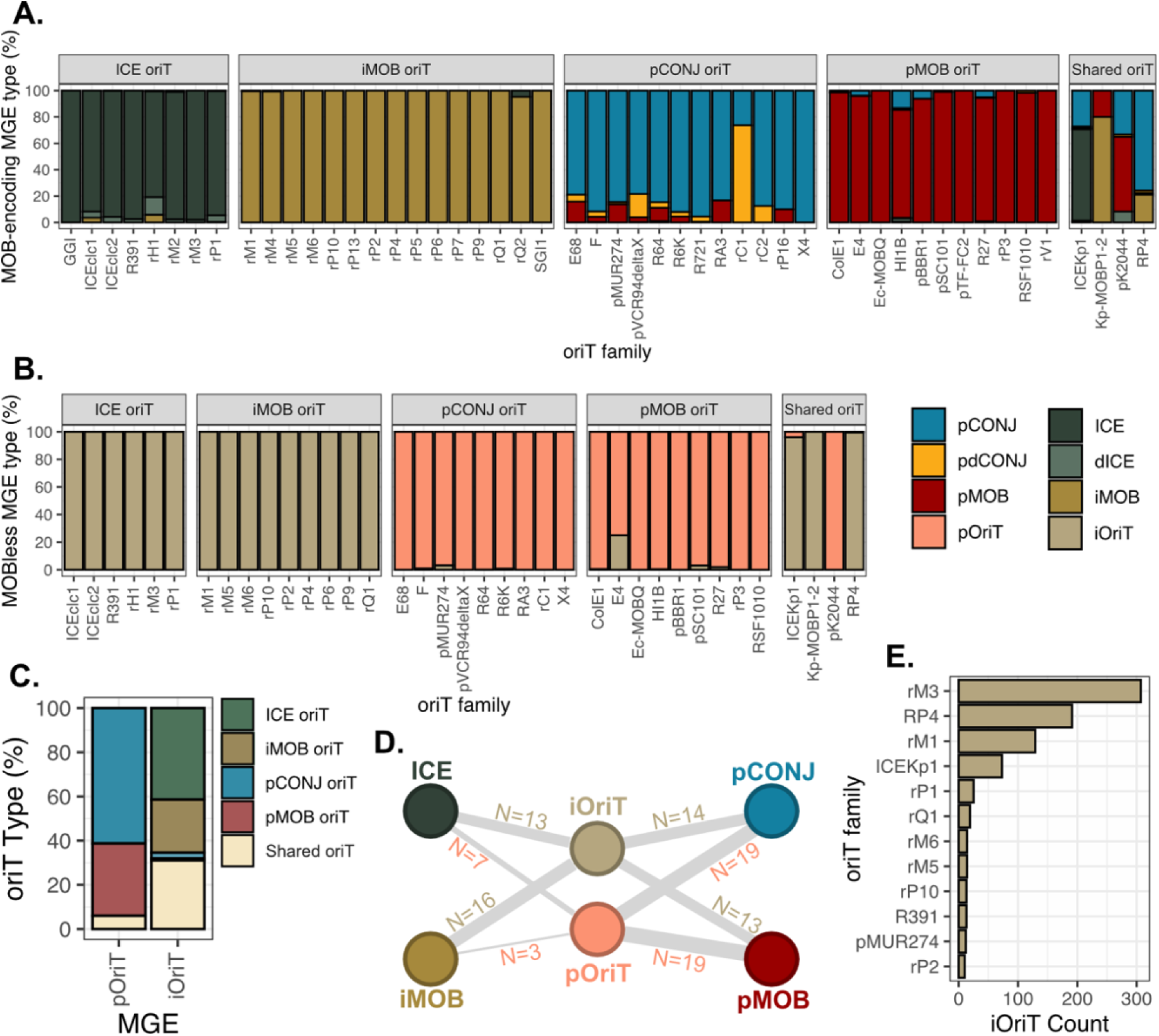
**A.** Frequency of MGE types carrying a relaxase in which an *oriT* family has been identified. The *oriTs* were split into ‘ICE *oriT’*, ‘iMOB *oriT’*, ‘pCONJ *oriT’* and ‘pMOB *oriT’*, according to the most common MGE associated to the family. A fifth group ‘Shared *oriT’* contained those associated to different MGEs. Only *oriT* families with 10 or more occurrences in the dataset are shown. **B.** Frequency of MOBless MGEs in which an *oriT* has been identified. Only *oriT* families with 10 or more occurrences in the dataset are shown. The *oriTs* were grouped into the same five categories as 4A. **C.** Percentage of *oriT* categories carried by pOriTs and iOriTs. **D.** Graph showing the number of *oriT* families with >10 occurrences shared between MOBless elements and pCONJ, pMOB, ICE and iMOB. The nodes of the graph represent the MGEs. Edges connect MGE types when a same *oriT* has been found in both elements, its width representing the number of different *oriTs* in common. The number is shown next to the edge. **E.** Number of *oriT* families associated to iOriTs. Only cases with >10 iOriTs are shown.

The observation that plasmid *oriT*s are rarely found in chromosomes (Fig 2E, Fig S4), suggests a split of *oriT* families between integrative elements and plasmids. Indeed, the vast majority of *oriT* families of elements encoding a relaxase are specialized: either they are found in plasmids or in integrative elements (Fig 4A and 4D). The frequency of iOriT and pOriT in each *oriT* family confirms the rule that *oriT* families tend to be specialized in terms of the type of MGE (integrative or plasmid), most exceptions caused by very low counts for these *oriTs* in MOBless elements (Fig 4E). Only few *oriT* families are found at appreciable frequency in both types of elements (coined “*shared oriTs*” in Fig 4A-C) and only two are frequent (Fig 4F): *oriT*_RP4_ which is found in many pCONJ and iMOB, and *oriT*_ICEKp1_ which is found in many ICEs and pCONJ (Fig 4A). Interestingly, they are also found in many MOBless elements, but almost exclusively in iOriT (nearly absent in pOriT), suggesting complex interactions between some GIs and plasmids.

To detail the interdependencies between different MGE types, we retrieved all *oriT* families with >10 occurrences in our dataset and built a graph with the number of *oriT* families than can be found in two different types of MGEs. Our results show that plasmids are highly interconnected, *i.e.,* pOriTs heavily rely on other plasmids for their mobility (Fig 4D, Fig S12). In contrast, iOriTs have *oriT*s from families present in ICEs and pCONJ (Fig 4D). Among the iOriTs, 335 have an ICE-like *oriT*, 205 have an iMOB-like *oriT*, 265 have *oriT*s of families with multiple origins (“*shared oriTs*”), and only 30 have a plasmid-like *oriT*. This result suggests that ICEs and IMEs drive the transfer of most Genomic Islands lacking relaxases, but some conjugative plasmids may also contribute to their mobility. In contrast, interactions between iOriT and pMOB or between pOriT and iMOB can occur (Fig 4D), but are rare (Fig 4C).

### iOriTs form large families of MGEs

The above data suggests that iOriTs are specific families of MGEs with importance in bacterial evolution. Alternatively, iOriTs might just be decaying MGEs that recently lost conjugation-associated protein coding genes. If so, then iOriTs should be intermingled in families of ICEs and IMEs. To test this hypothesis, we computed the weighed Genetic Repertoire Relatedness (wGRR, see Methods) between the 4,298 *att*-delimited ICEs and IMEs and the 10,492 conjugative and mobilizable plasmids (discarding 773 phage-plasmids and 1,825 non-transmissible ones) (Fig S13). Briefly, wGRR=1 means two elements have identical genetic repertoires, or one element is a subset of the other, as expected if one is decaying, whereas wGRR=0 indicates lack of homologous genes. We found 590 elements with no wGRR hit with other element in the dataset. The remaining 14,200 elements were clustered into 438 communities using the Louvain algorithm on the wGRR matrix (see Methods). Since most communities had few elements, we retrieved the 181 communities with more than 5 elements for further analysis (including 13,442 elements) (Fig S13). In most of these communities, more than 75% of the elements are of the same MGE type (N=161/181). The 723 iOriTs were found in 41 communities, of which 21 iOriTs were in those dominated by other MGEs (mostly iMOBs). A set of 87 iOriTs accounted for ~50% of the elements of eight mixed communities, in which case the whole community typically shared the same *oriTs* family. These cases of iOriTs highly similar to iMOBs could be due to recent events of MOB loss in the former. In contrast, 615 iOriTs constituted 19 communities where they were the only or the most numerous element (Fig 5A, Fig S14). Among these communities, four had elements from two or three different species (Fig S15), showing inter-species dissemination of highly related iOriTs. These results suggest that while some iOriT may have recently derived from iMOBs, most iOriTs are in large families of similar types of elements.

**Figure 5.**
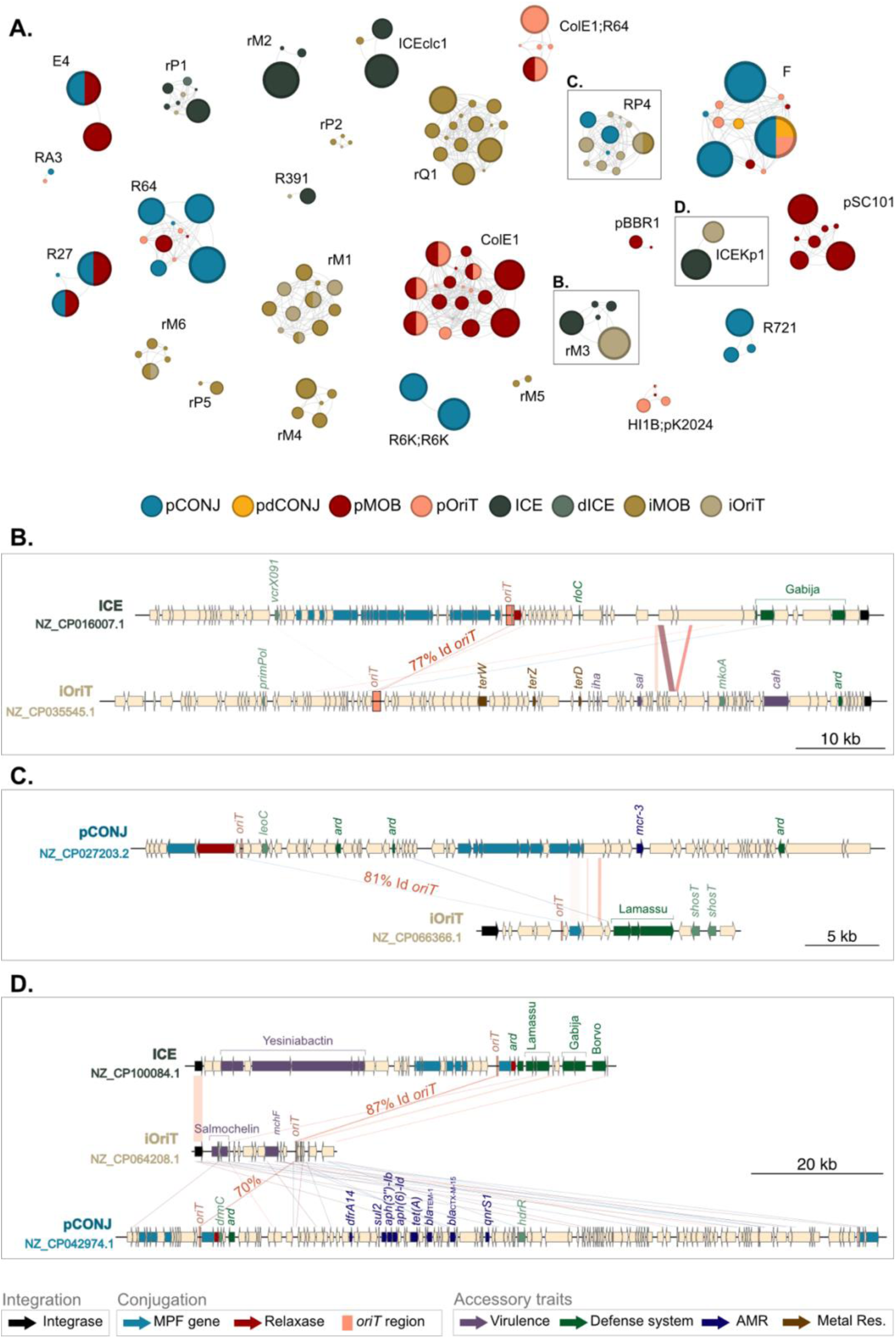
**A.** MGE communities carrying the same *oriT* families. Nodes represent communities, their size indicating the community size (number of elements) and color indicating the dominant MGE (>75% of the community). When more than one color is shown, it indicates two frequent MGEs. Nodes are connected when communities carry the same *oriT* family, its name being indicated next to the cluster. The three grey squares highlight the communities further detailed in B., C., and D., respectively. **B-D.** Sequence alignment of elements carrying the *oriT* families *oriT*_rM3_, oriT_RP4_ and *oriT*_ICEKp1_, respectively. The arrows indicate genes and their colors the functional groups (legend at the bottom).

To explore the functional dependencies of iOriTs, we built a graph in which each node represents a community and edges connect communities with the same origin of transfer (Fig 5A). This revealed pairs of families of putative hitcher and helpers that have similar *oriTs* but very different genetic repertoires (e.g., see clusters with *oriTs* rM3, RP4, ICEKp1) (Fig 5A). For instance, iOriTs with the *oriT*_rM6_, are the TAI islands identified in *E. coli* O157:H7 in 2002, whose transfer mechanism remained unknown until now^55^. Remarkably, their *oriT* region is 60% identical to the one recently reported in the ICE SGI-4^56^, whose MOB_M_ relaxase had not been identified prior to this study. These iOriTs have almost no sequence similarity to the ICEs able to mobilize them beyond the *oriT* region (Fig 5B). In other cases, the hitcher-helper pairs are very different in size and may even differ in terms of vertical transmission (some are integrative, and the others are plasmids). For instance, different families of small iOriTs and iMOBs carry the same *oriT* family as the well-known conjugative plasmid RP4 (Fig 5C), and have been simultaneously found across other Pseudomonodota^57^. This is often the only region of similarity between the different types of elements. Lastly, there are iOriTs with *oriT* families found in both ICEs and pCONJ, which suggests they may exploit one or both types of elements. This is the case of small iOriTs carrying the *oriT*_ICEKp1_ also found in the ICE*Kp1* described in *K. pneumoniae* in 2008^58^ (Fig 5D), and also carried by conjugative plasmids (Plasmid Taxonomic Unit PTU-E50) (Fig 5C). None of the three groups of elements shows high sequence similarity beyond the *oriT*s.

### iOriTs provide critical traits to their hosts

Given the diversity of sizes and gene repertoires of iOriT (Fig 5B-D), we assessed their potential functional contribution to bacterial adaptation. As is typical of MGEs, the most common categories are unknown functions (S) or replication-recombination (L) (Fig S16). However, iOriTs were enriched in specific categories compared to other elements, such as signal transduction mechanisms (T) or amino acid metabolism (E). We then focused on specific functions often associated with genomic islands in the literature^28^, including virulence factors, antibiotic resistance genes, and anti-MGE defense systems (Fig 6A). These functional repertoires vary across types of MGEs, with the density of virulence factors being higher in iOriT than in any other types of integrative elements or plasmids (Fig 6A, Fig S17). Defense systems are present in all MGEs, while counter-defense tend to be less frequent in iOriTs (Fig S16). While antimicrobial resistance is very frequent in pOriT^6^, it is almost absent from integrative IMEs (Fig 6A). This is intriguing as some iOriT carry *oriT*_ICEKp1_ that are also carried by plasmids with antimicrobial resistance genes (Fig 5C). In contrast, many iOriT encode tellurite resistance, which is their only common resistance trait (Fig 6D). The reasons for these differences are intriguing and suggest that iOriT impact some processes (virulence, defense, metal resistance) more than others (antibiotic resistance).

**Figure 6.**
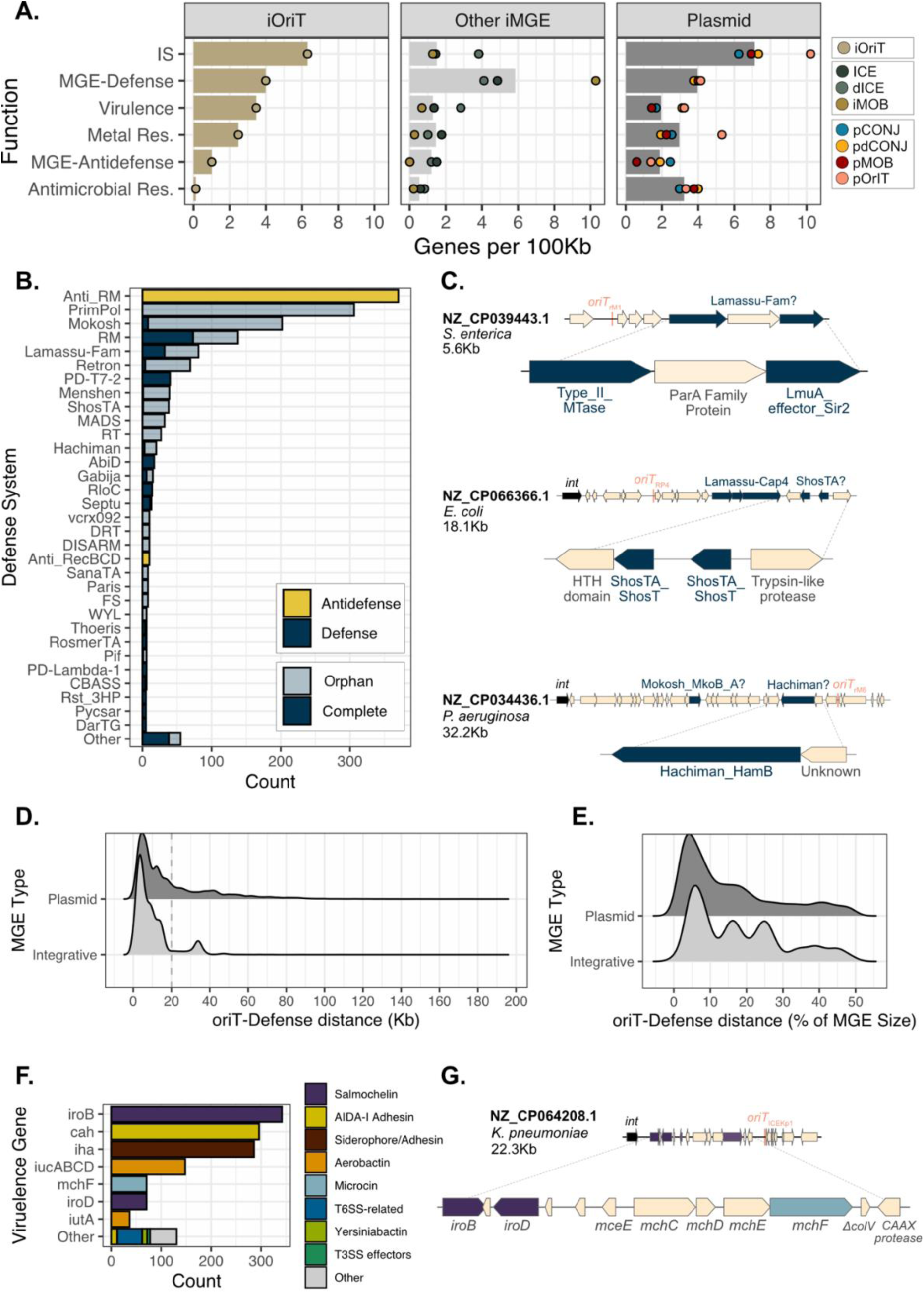
**A.** Gene density (gene number per 100Kb) of selected traits/functions encoded in iOriTs (left), other integrative MGEs (center), and plasmids (right). The colored dots indicate the gene density for each MGE type. **B.** Most common defense systems (blue) and antidefense genes (yellow) identified across iOriTs. Orphan genes (genes identified in the absence of a complete system) are shown in a lighter blue. **C.** Three cases of orphan genes in iOriTs. In each case, the full iOriT is represented next to the accession number of its genome, species, and size (upper part), and the location and annotation of its defense and adjacent genes (bottom). **D.** Distribution of the absolute distance (Kb) between every defense system and anti-defense gene and the origin of transfer. **E.** Distribution of the distance between the defense and anti-defense systems compared to the MGE size (percentage of its genome size). **F.** Most common virulence genes identified across iOriTs. **G.** Genetic map of an iOriT with the *oriT*_ICEKp1_. The virulence genes are highlighted, their color matching the legend of panel D.

We identified many MGE-defense and antidefense genes in iOriTs (5.01 genes/100Kb), in agreement with recent studies in other MGEs^59^. Among these, genes encoding anti-restriction enzymes, restriction-modification systems, and orphan methylases are the most common (Fig 6B). Anti-restriction proteins were found on large iOriTs and are nearly absent in smaller elements (Fig S18). Similar trends were observed in plasmids and could be explained by smaller elements bypassing restriction barriers without the need of expressing anti-restriction^60^. Anti-MGE systems frequent in iOriT include Lamassu-like systems, which were proposed to contribute to the rarity of plasmids in the 7^th^ pandemic *V. cholerae* strains^61^. Lamassu is the 3^rd^ most common complete defense system in iOriTs, with more than 500 occurrences across ICE, dICEs, and iMOBs. Remarkably, we found 956 genes predicted as being part of defense systems but lacking the remaining genes. They are often co-localized with unknown function genes or other complete defense systems (Fig 6C), suggesting they could be novel defense systems (or variants of known ones). Most defense systems found in iOriTs and other ICEs and IMEs were located close to the origin of transfer (Fig 6D-E, Fig S19, see Methods). Next to the *oriT* lies the leading sequence that first enters the recipient cell during conjugation^62^. It was recently shown that leading sequences of conjugative plasmids contain multiple anti-defense systems allowing the plasmid to counteract the recipient cell defense systems^63^. These results suggest that integrative elements transferring by conjugation also use anti-defense and defense systems in leading sequences to increase their chances of successful transfer.

Strikingly, many iOriTs encode virulence factors, having a much higher density of these genes (3.47 genes/100Kb) than ICEs. The iOriT communities include known Pathogenicity Islands (Fig 6F). For instance, the large family of iOriT with *oriT*_M5/6_ previously identified as TAI islands encode the virulence *iha* gene which encodes for an adhesin important in urinary tract infections (Fig 5B)^55^. The community of small iOriTs carrying the *oriT*_ICEKp1_ (previously identified as GIE492) encodes the microcin and salmochelin genes associated to hypervirulent *K. pneumoniae* (Fig 6G)^64^. Crucially, they have *oriTs* similar to those of the known pathogenicity island ICE*kp1* and often co-occur in the same genomes (Fig S20), suggesting that elements of different type may be co-mobilized together to provide a larger array of virulence factors to pathogenic bacteria.

## DISCUSSION

Bacteria harbor a plethora of Genomic Islands acquired by Horizontal Gene Transfer, but whose mobility mechanisms remain elusive. Here, we further developed our method to identify novel *oriT* families to find thousands of integrative elements mobilizable by conjugation, including over 1,500 putative iOriTs. This suggests that many GIs can be mobilized by conjugation via *oriT* mimicry. We have reasons to believe this large number of integrative hitchers is still an underestimate. Even if we could identify *oriT*s in almost all ICEs, we may have failed to identify some *oriTs,* especially those encoded far from the relaxases or near previously unrecognized relaxases. The latter should not be underestimated, as revealed by our observation that numerous MOB_M_ relaxases were previously undetected among some of the best studied bacteria. Lastly, hitchers may use mechanisms that we have not studied here. For example, MGIVchHai6 uses a relaxosome accessory protein (MobI) to exploit conjugative plasmids by modifying the affinity of their relaxases, thereby having a different *oriT* from the ones contiguous to the helper’s relaxase^65^. Some hitchers integrate in or next to ICEs^66^, or in conjugative plasmids^67^, to transfer by conduction, which dispenses them from encoding an *oriT*. Some IMEs integrate ICEs and their excision is necessary to the transfer of both elements^68^. They produce complex integrative elements that may be difficult to disentangle without dedicated experimental studies. Finally, ICEs may conjugate before excision from the chromosome, thereby transferring neighboring chromosomal sequences and MGEs^50,69,70^. Here, we showed that some iOriTs are previously identified Pathogenicity or Defense Islands whose mechanisms of mobility remained unknown, sometimes decades after their discovery. Further work is likely to reveal an even greater number of conjugative hitchers.

This work reveals that integrative elements mobilizable by conjugation are much more abundant than ICEs (Fig 3A). Previous works showed fewer differences^14,16^, which is explained by the identification of the iOriT that is novel to this work and by more sensitive identification of the very abundant MOB_M_ relaxases. A large excess of integrative hitchers relative to conjugative helpers was also previously observed in plasmids^8^. Together, these works show that most elements being mobilized by conjugation are not autonomous and require the help of a conjugative element.

Intriguingly, the frequency of each type of MGE largely depends on the species: integrative MGEs largely outnumber plasmids in *P. aeruginosa, S. enterica* and *V. cholerae* and are clearly outnumbered by the latter in the other three species (Fig 3B). This wide variation could partly be explained by the presence of specific anti-MGE systems, like Lamassu. While only a few specifically anti-plasmid systems have been uncovered^61,71,72^, the current focus on anti-phage systems may have hidden a larger diversity of mechanisms that could be present among the many chimerical or putatively partial defense systems uncovered by ours and others analyses (Fig 6BC). Anti-plasmid systems encoded by integrative MGEs may thus directly contribute to the rarity of plasmids in some genetic backgrounds. In addition, intrinsic properties of the two different types of elements may contribute to the observed differences in their frequency. For example, it was proposed that conjugative plasmids tend to evolve faster in terms of gene repertoires, whereas ICEs may be more stable and have broader host range^46^. It would be important to test if such differences also apply to their hitchers.

The identification of *att* sites in many elements allowed for the first time to systematically query the size and gene repertoires of integrative elements at a large scale. While our methods differ, we found an excellent concordance with the much broader data on Genomic Islands by TIGER^47^, which did not specifically searched to characterize hitchers. The clustering of integrative elements revealed clearly separated groups that tend to be homogeneous in terms of genetic mobility, i.e. that are dominated by one type (like the iOriT groups in Fig 5). This suggests that differentiation between elements transferred by conjugation is ancient, although transitions between types mediated by deletions can be observed. Recently, there have been attempts to class plasmids using genetic similarity into groups, notably in Plasmid Taxonomic Units^73^. Our results now suggest that such methods might be used with equal success to class integrative elements. Since MGEs often cross the boundaries of our definitions^42^, we propose to call them Conjugative Taxonomic Units by analogy. As a rule, our results show that within a group all or most elements have the same *oriT* family.

Hitchers are expected to exploit helpers with *oriT*s of the same family, even though we cannot know at this stage if a hitcher can exploit many or just a few of the potential helpers carrying the same type of *oriT*. Nevertheless, an intriguing finding of our work is the specificity of MGEs relying on *oriT* mimicry: plasmid hitchers tend to exploit conjugative plasmids, while integrative hitchers tend to exploit ICEs (Fig 4). This specificity should not derive from fundamental differences in the conjugation mechanisms, which are very similar and share a long evolutionary history^13,74^. Instead, the reasons of this specificity could be explained by the processes of emergence of hitchers. Firstly, hitchers may evolve from helpers by total or partial loss of the MPF locus, as previously found in plasmids ^49^. This results in hitchers and helpers having similar *oriT*s. This is consistent with the observation of 21 iOriTs in communities dominated by other types of elements. Secondly, functional dependencies between integrative and plasmid elements may require additional layers of genetic regulation. Conjugation of integrative elements requires the previous excision and circularization of the element to avoid conjugation of large fractions of the bacterial chromosome^75,76^. An IME acquiring an *oriT* from a conjugative plasmid would thus be at the risk of constantly conjugating the chromosome. Accordingly, excision and conjugation were found to be tightly regulated in ICEs and IMEs^77–80^. Some hitchers do not follow this rule. The iOriTs carrying an origin of transfer similar to that of the conjugative plasmid RP4 form large communities across several species suggesting they are ancient iOriTs with gene repertoires unrelated to those of RP4-like plasmids. Indeed, a parallel study has simultaneously revealed the presence of these IMEs carrying *oriT*_RP4_ across 100 species of Pseudomonodota^57^. These prevalent integrative elements seem to have evolved the ability to hijack these conjugative plasmids *de novo*, e.g. by acquiring the *oriT* by genetic exchanges. Exchanges between different types of MGEs, including ICEs and plasmids, have described numerous times^81,82^ and could also contribute to the emergence of hitchers.

Conjugation can transfer huge amounts of DNA in one event, up to a complete genome^83^, contrary to other mechanisms like transduction^84,85^. Yet, there are very specific size distributions for these elements: ICEs tend to be large, iMOBs small, and the distribution of sizes of iOriT is bimodal. Analogous plasmids follow similar distributions, with the difference that pMOB plasmids are also largely bimodal^6^. The differences in size between ICEs and IMEs was previously identified and is likely due to the requirement of encoding a MPF which can require from 11 to dozens of genes^36^. While plasmids and integrative elements have approximately the same average size, their ranges of variation are different. The smallest IMEs are larger than the smallest mobilizable plasmids, possibly because of the sequence required to encode the integration genetic module and its regulatory components. Indeed, while more than 1,200 plasmids (11.5%) encode less than 5 genes, only 12 IMEs (0.5%) were in the same situation (Fig S21). At the other edge of the distribution, while large ICEs have been described (e.g. the 800 kb tri-partite ICE^22^) there are many more very large plasmids in our dataset. Larger integrative elements might be costly by disrupting chromosomal organization. In the other side, smaller integrative elements might result from gene deletion *in situ*, as observed by the higher number of decaying elements in chromosomes (dICEs) compared to plasmids (pdCONJ) (Fig 1E). Decaying pdCONJ are rapidly lost^49^. In contrast, dICEs are not affected by the segregationally loss imposed to plasmids and may remain in genomes for longer periods of time.

An intriguing observation of this study is that among elements mobilizable by conjugation, those that integrate the chromosome encode traits very different from those that are plasmids. While pOriT are enriched in antimicrobial resistance, IMEs are enriched in MGE-defense systems, and iOriTs are specifically enriched in virulence factors. Antimicrobial resistance has been associated to conjugative, fast evolving plasmids^86^, while ICEs were proposed to be less plastic^46^. This might explain the differences we observe. Of note, integrative elements have been associated to AMR in other species^16,87^, and it’s possible that such differences are species specific. In contrast, iOriT encode many anti-MGE systems and a high density of virulence factors. This raises an interesting parallel between MGEs mobilized by conjugation and temperate phages, whose integrative elements (prophages) and their hitchers (phage satellites) also often encode virulence factors and defense systems, and rarely encode antibiotic resistance^59,88^, whereas phage-plasmids encode more frequently antibiotic resistance^89^. Further work is needed to understand these striking, but poorly understood, associations between the mechanisms of vertical and horizontal transmission of MGEs and the accessory traits they encode.

## MATERIALS AND METHODS

### Genome data

We retrieved the 32,799 complete bacterial genomes from the NCBI non-redundant RefSeq database (https://ftp.ncbi.nlm.nih.gov/genomes/refseq/, accession in May 2023). Only the 6,182 genomes belonging to *Escherichia coli* (n=2,559), *Salmonella enterica* (n=1,293), *Klebsiella pneumoniae* (n=1,522), *Vibrio cholerae* (n=107), *Neisseria gonorrhoeae* (n=142), and *Pseudomonas aeruginosa* (n=559), were selected for further analysis. These genomes had one or two chromosomes (*V. cholerae*) and 13,090 plasmids: *Escherichia coli* (n=6,135), *Salmonella enterica* (n=1,465), *Klebsiella pneumoniae* (n=5,154), *Vibrio cholerae* (n=16), *Neisseria gonorrhoeae* (n=166), and *Pseudomonas aeruginosa* (n=154). Accession numbers of genomes can be found in Table S2.

### Functional annotation

Genomes were annotated using Prodigal v2-6-3, with the option *-p meta*, recommended for mobile genetic elements^90^. The COG categories of the proteins were inferred using the software eggNOG-mapper, version 2.1.12^91^. We used the module CONJscan of MacSyFinder, version 2.0 to identify protein coding genes related to conjugation, such as relaxases and the MPF systems^92^. MGE defense systems were screened using DefenseFinder, v1.3.0, retrieving both complete systems and orphan hits using the option *--preserve-raw* ^93^. Antidefense systems were also screened using the option –*antidefensefinder*^94^. Genes related to antimicrobial resistance, heavy metal resistance, stress response, and virulence were identified with AMRFinderPlus, version 3.11.4^95^. Virulence genes were additionally screened using BLASTp of BLAST 2.9.0+^96^, using as query the 27,982 protein sequences available in the virulence factor database VFDB (http://www.mgc.ac.cn/VFs/, accession in August 2024)^97^, retrieving only the hits >50% identity and >50% coverage. Insertion Sequences were screened using ISEScan, version 1.7.3^98^, with default parameters. Integrases were identified screening the PFAM domains for tyrosine recombinases (PF00589.27) and serine recombinases (PF00239.26 and PF07508.18), with a coverage >40%^99^. For serine recombinases, only proteins containing both domains were considered.

### Identification of occurrences of origins of transfer of known families

*oriTs* of known families were retrieved from three different collections: an in-house collection of 96 experimentally validated *oriTs* from the literature^6^; a collection of 122 experimentally validated *oriTs* available in the updated database oriTDB^100^; and 21 *oriT*-comprising regions recently inferred from *E. coli*, *K. pneumoniae* and *Acinetobacter baumannii*, six of which were experimentally validated^35^. To note, an updated sequence of the region *oriT*_FK_ and extended versions of *oriT* ^101^ and *oriT* ^102^ are provided. After removing redundant *oriTs* across the collections, we obtained a final dataset of 168 non-redundant known *oriTs* and *oriT*-comprising regions (Dataset S1). Known *oriTs* were clustered by sequence similarity into *oriT* families. This was done using the Smith-Waterman algorithm of the FASTA package, v36.3.8i (ssearch36, E-value<0.001)^103^. The parameter we used for the clustering is the alignment bitscore since it is independent of the database size and directly reflects the sequence similarity and alignment length. However, the bitscore depends on the pair of aligned sequences, and cannot be used directly to compare different alignments. Instead, it can be normalized in a value between 0 and 1 by comparing how well two sequences align to each other, relative to how well they align to themselves (i.e., the maximum bitscore for that particular pair). This ensures a length-independent, symmetric, and comparable normalized bitscore. To do so, we employed the following formula:

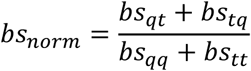

where *bs_norm_* is the normalised bitscore; *bs_qt_* is the bitscore of the alignment query vs. target; *bs_tq_* from target vs. query; *bs_qq_*, query vs. query; and *bs_tt_*, target vs. target. The *bs_norm_* was used for hierarchical clustering analysis of the non-coding sequences using the Ward.D2 method of the R package pheatmap (v.1.0.12) (https://CRAN.R-project.org/package=pheatmap). The 168 *oriTs* were grouped into 90 *oriT* families, named after well-known *oriTs* they contain (Table S3). For instance, the family *oriT*_F_ includes ten non-redundant *oriTs* from the three sources: *oriT*_F_, *oriT*_ColB4_, *oriT*_P307_, *oriT*_pSU233_, *oriT*_R1_, and *oriT*_R100_ of our in-house collection; *oriT*_F_, *oriT*_pEC156_1_, and *oriT*_pKPHS2_ of the oriTDB; and the *oriT*_FK_ containing region. Known *oriT* families were screened using BLASTn of BLAST 2.9.0+ ^96^, with the options *-task blastn-short -evalue 0.01*. Only the hits with an identity >50% and coverage >50% were retrieved.

### Identification of previously uncharacterized origins of transfer

To identify unknown *oriT*-comprising regions across the complete genomes, we used a method recently described for plasmids^35^. Briefly, we retrieved the first intergenic sequence >50nt upstream the relaxase genes of both chromosomes and plasmids. Among the 14,429 sequences, redundant sequences were identified clustering by sequence similarity using MMSeqs2, version 13.451^104^, with a criterion of 100% identity and 100% coverage. Only one representative per cluster was randomly picked for subsequent analysis. This resulted in a dataset of 3,111 non-redundant intergenic sequences upstream a relaxase that were split into seven groups according to their MOB family: 242 MOB_C_, 373 MOB_F_, 461 MOB_H_, 186 MOB_M_, 1255 MOB_P1_, 408 MOB_Q_ and 70 MOB_V_. Per group, we performed an all-vs-all local alignment of the intergenic sequences using the Smith-Waterman algorithm of the FASTA package, v36.3.8i (command ssearch36)^103^, retrieving only the hits with an E-value <0.001, identity >50%, coverage >30%, alignment length >50 nt.

The results were used to cluster the sequences using the Ward.D2 method of the R package pheatmap, version 1.0.12 (https://CRAN.R-project.org/package=pheatmap), based on the normalized bitscore (see above). This clustering was performed for each MOB family independently. Hierarchical clustering may encounter difficulties when aiming at unveiling small clusters in the presence of large ones. To tackle this limitation, we performed the clusters in serial rounds. For instance, in case of MOB_P1_, we clustered the 1,255 sequences into the 17 largest clusters (average 57 sequences per cluster), leaving the remaining 288 sequences as an outgroup. In a second round, the outgroup was clustered into 57 clusters (4 per cluster). Small clusters (<5 sequences/cluster) were discarded, resulting in a total number of 36 clusters. Each cluster represents a candidate *oriT*-comprising region for MOB_P1_. The same procedure was followed for each MOB group, resulting in 212 clusters (2,417 sequences). Among these *oriT*-comprising regions, 29 were discarded since they belong to plasmids where the *oriT* is not located upstream to the MOB (e.g., *oriT*_F_, *oriT*_pVCR94X_).

We performed different validations with the remaining 183 clusters. First, a multiple sequence alignment (MSA) was performed to validate the quality of the clusters. We used mafft, version v7.505, with the options *--maxiterate 1000 –genafpair*^105^. The MSAs were then examined with the software SnapGene, version 7.0.3 (www.snapgene.com), and 11 clusters were discarded due to the bad quality of the MSA. Second, we examined the genetic location and surrounding areas of the sequences using SnapGene, version 7.0.3 (www.snapgene.com). Among them, 41 clusters were discarded due to either gene rearrangements (e.g., Insertion Sequences upstream the relaxases), or wrong annotations (e.g., the first intergenic sequence was not properly annotated). Among the 131 remaining clusters, some were grouped into the same *oriT* family since their sequences contained the same known *oriT* (*e.g.*, *oriT*_R46_ was identified across the sequences of two clusters), finally reaching 115 clusters. After the first screening (see below), these 115 *oriT*-comprising regions had recognizable instances in the 87% of the relaxase-carrying Genomic Islands. For the sake of completeness, we repeated the same pipeline with the relaxases lacking *oriTs* without removing redundant intergenic sequence prior to the hierarchical clustering. The analysis resulted in 9 additional clusters, leaving only 7% of the relaxases without a putative *oriT*.

These 124 *oriT*-comprising regions were clustered, together with the previously known *oriTs*, by sequence similarity into *oriT* families, as previously explained. This resulted in a total of 76 families of origins of transfer, of which 52 are new. Novel *oriT* family regions were named according to their MOB associated and a numeric identifier. To distinguish these *oriT*-comprising regions from delimited *oriT* sequences, an ‘*r’* standing for ‘*region’* was added to the name. Therefore, the families rQ1 (*oriT*_rQ1_) and rQ2 (*oriT* _rQ2_) correspond to two different *oriT*-comprising regions associated to MOB_Q_ relaxases.

### Identification of *oriTs* distant to the relaxases

Since the HMM profiles used to screen for *oriTs* were built using the clustering of intergenic sequences upstream relaxases, our approach misses those few *oriTs* known to be located far from the relaxase: *oriT*_F_^3^, *oriT*_pVCR94X_^106^, *oriT*_R391_^107^ *oriT*_ICEclc2_^45^ *oriT*_pK2044_^108^. So, we retrieved all the blastn hits obtained from each of these queries (as previously explained in ‘Screening of known origins of transfer’) with the tools from SeqKit, version 2.8.2^109^, to perform a MSA using mafft version v7.505^105^ and build HMM profiles using HMMER, version 3.3.2^110^.

### Screening of the uncharacterized *oriTs* across the complete genomes

To screen for the candidate *oriT*-comprising region across the complete genomes, we used the MSAs previously performed to build Hidden-Markov model (HMM) profiles of each of the clusters. First, the MSAs were manually trimmed in SnapGene, version 7.0.3 (www.snapgene.com). The edges were removed, keeping the alignment from the first ten consecutive nucleotides in a row within the consensus sequence to the last ten consecutive nucleotides in a row within the consensus sequence. Additionally, every sequence whose length was <50% or >50% of the consensus sequence of the trimmed MSA was discarded. In cases where an already known *oriT* was identified in the *oriT* regions, the former was included in the MSA, and the trimming was performed from the beginning to the end of the experimentally validated origin of transfer. The trimmed MSAs were then used to build HMM profiles employing the command hmmbuild of HMMER, version 3.3.2^110^. Once the HMM profiles were built, we used nhmmscan to screen for the *oriTs* and *oriT*-comprising sequences across the 6,182 complete genomes.

### Validation of the *oriT* diversity within the species

To assess if the novel *oriTs* inferred in this work cover the *oriT* diversity within the species, we retrieved a supplementary dataset including the 2,289 genomes of the six species submitted between June 2023 and May 2025 (https://ftp.ncbi.nlm.nih.gov/genomes/refseq/, accession in June 2025). The genomes were screened for relaxases and *oriTs* as previously described.

### Identification of defense and anti-defense systems in the conjugative leading region

Defense/anti-defense systems were searched in the first DNA region to enter the cell during conjugation (i.e, the conjugative leading region), defined in plasmids as the one adjacent to the *oriT* in the opposite direction to the relaxase and/or relaxosome accessory protein genes (Fig S19A)^2,3,63^. Most conjugative plasmids in these species carry a MOB_P1_ or MOB_F_ with the *oriT* locating at one side of the relaxase-MPF module (Fig S19B)^3^. In these cases, the leading region is in accordance with the previous definition. However, that in many integrative elements, such as definition was not necessarily warranted. The conjugative machineries with MOB_C_ or MOB_M_, which are the most abundant in ICEs (and also present in plasmids, see below), harbor the *oriT* between the MPF operon and the relaxase gene (Fig S19C)^58^. Even more complex scenarios arise with MOB_H_ systems, where the *oriT*, relaxase, and different modules of the MPF system are encoded in different regions of plasmids^111,112^, thereby complicating the *in silico* identification of the leading region (Fig S19C). Given the diversity of such cases and in lack of experimental data for these elements, we computed the absolute distance between the *oriT* and every anti-defense gene and defense system identified with DefenseFinder v1.3.0^94^. Only elements with one *oriT* were considered for the analysis.

### Classification of Mobile Genetic Elements

Integrative MGEs were classified according to their mobility, following the most recent classification of plasmids (Fig 1A)^42^. Briefly, MGEs encoding a relaxase and a complete MPF system were classified as Integrative Conjugative Elements (ICEs) when integrated in the chromosome, or conjugative plasmids (pCONJ) when episomal. MGEs encoding an incomplete MPF system, with or without a relaxase, were considered decayed ICEs (dICEs) or decayed pCONJ (pdCONJ). Integrative elements missing a MPF system were classed as Integrative Mobilizable Elements (IMEs) and could carry their own relaxase (iMOB) or only an *oriT* (iOriT). Likewise, plasmids carrying an *oriT*, with or without a relaxase gene, were classed as mobilizable plasmids (pMOB and pOriT, respectively). Plasmids with phage-like functions characteristic of phage-plasmids were identified using the tyPPing pipeline^113^. The remaining plasmids were considered non-transmissible (pNTs). Plasmids were assigned to a Plasmid Taxonomic Unit (PTU) when possible, using COPLA, version 1.0^114^. By using the module CONJscan of MacSyfinder, version 2.0^92^, plasmids, ICEs and IMEs encoding a relaxase were assigned to one of the nine MOB classes encompassing the phylogenetic diversity of relaxases^115^: related to HUH-endonucleases (MOB_B_, MOB_F_, MOB_P_, MOB_Q_, MOB_V_), HD-hydrolases (MOB_H_), restriction endonucleases (MOB_C_), rolling-circle replicases (MOB_T_), or related to tyrosine recombinases (MOB_M_). Also, elements carrying a complete or incomplete MPF system where assigned to one of the eight MPF monophyletic groups^13^: MPF_I_, MPF_C_, MPF_G_, MPF_T_, MPF_F_, MPF_B_, MPF_FA_, or MPF_FATA_. Detailed information on the plasmids and integrative elements is available in Supplementary Tables S4-S6.

### Identification of MOB_M_ relaxases

Over the course of the project, we realized that the state-of-the-art methods to identify relaxases missed occurrences of the MOB_M_ family (not to be confused with the MobM protein of pMV158, which is MOB_V_ family^116^). Just three occurrences of the relaxase have been described in the literature: TcpM in the conjugative plasmid pCW3 of *Clostridium* (ABC96296.1)^40^, MobK in the mobilizable plasmid pIGRK of *Klebsiella* (AAS55463.1) ^39^, and MpsA in the IME SGI1 of *Salmonella* (AAK02039.1)^37^. The screening for MOB_M_ proteins in the available software relies on a HMM profile based on the *Clostridium* pCW3 relaxase and it overlooks phylogenetically distant MOB_M_. Given the species composition of our database, we built a new HMM profile for their identification.

First, we identified MOB_M_ in our dataset using two rounds of BLASTp of BLAST 2.9.0+^96^. The first round used seven relaxases as query - the three known MOB_M_, and four other instances identified in our dataset (WP_033567157.1, WP_047656026.1, WP_001096463.1, WP_004252296.1). Only the hits with an E-value <0.001 and coverage >30% were retrieved. The 1,156 proteins identified in the first round were then used as query in a second BLASTp, retrieving 1,521 hits with an E-value <0.001, identity >30%, and coverage >50%. Across the latter proteins, identical sequences were deduplicated using MMSeqs2, v.13.4511^104^. The 143 deduplicated MOB_M_ sequences were aligned using MAFFT, version 7.505, options *--maxiterate 1000 –genafpair*^105^. The MSA was trimmed from the edges and used to build an HMM profile using hmmbuild from HMMER, v3.3.2^110^ (Dataset S2). Lastly, the HMM profile was used to screen for MOB_M_ relaxases across the dataset using hmmsearch and retrieving hits with an E-value <0.001. MOB_M_ relaxases are closely related to tyrosine recombinases^39^. Therefore, to avoid false positive results with these proteins, we set a score *cut_ga* of 84, which allows the identification of MOB_M_ relaxases while discarding tyrosine recombinases.

### Identification of interval-delimited Genomic Islands (GI^id^) across bacterial chromosomes

The genomic location of the MGEs were placed in relation to the location of persistent genes in each species to facilitate MGE delimitation and to compare locations across distinct genomes. For this, we computed the pangenome of each species using the pangenome module of Panacota, v1.4.1-dev3^117^, applying a minimum sequence identity threshold of 80% and a coverage threshold of 80%. We extracted from the pangenome the persistent gene families, i.e., those present in single copy in at least 90% of the genomes of the species using the module corepers of PanACoTA, v1.4.1-dev3^117^. We then defined *intervals* as the regions between two consecutive persistent genes in genomes. By definition, MGEs are not part of the persistent genome in most species. Hence, integrative MGEs are absent from persistent genes and constrained within intervals, providing upper bound to the position of the integrative MGES. We therefore defined interval-delimited Genomic Islands (GI^id^) as the interval where the MGE is present.

### Identification of genomic spots

Intervals from genomes of the same species were grouped in genomic spots when they are flanked by the same pair of persistent gene families, allowing comparison of equivalent regions across genomes. The number of genetic spots occupied by ICE/IMEs was generally low in the species (from 3 spots in *Ng* to 72 in *Pa*). Some intervals are flanked by unusual pairs of persistent genes, i.e. pairs that are rarely contiguous in the species. These usually result from the loss of one or both persistent genes or from the action of chromosomal rearrangements. These unusual configurations make the spot assignment ambiguous and MGEs in these regions were often subject to structural changes. Therefore, these intervals were excluded from the analysis (9% of all intervals).

### Identification of *att* sites and definition of *att-*delimited Genomic Islands

Since most ICE and IME integrate the chromosome using a Tyrosine recombinase, we used this information to propose a finer delimitation of the integrative elements. Integration sites were defined as follows: *attB* is the host attachment site prior to MGE integration (ancestral, MGE-free sequence), while *attL* and *attR* are the corresponding sites at the left and right boundaries of the integrated MGE. The latter share sequence similarity since a part of *attB* is duplicated and repeated at the edges of the MGE upon integration^118^. This allows their identification by searching for nucleotide sequence similarity within the spot. For each spot, we extracted the shortest interval (in bp) and assumed it represents the ancestral state containing the *attB* site (MGE-free) (Fig S9A). When this assumption does not hold we cannot identify the *att* sites. This interval was then compared with each longer interval containing an MGE. The longer intervals are expected to harbor short DNA repeats of the *attB* sites at the edges of the element, the *attL* and *attR* sites. To ensure at least two significant alignments per comparison enhancing detection reliability, all intervals were extracted including the persistent genes flanking them. To note, all intervals were oriented so that the flanking persistent genes are in the same direction, allowing to restrict the sequence similarity search to the positive strand. This improves computational efficiency and simplifies the identification of overlapping regions. Sequence similarity between the shortest interval containing the *attB* site (query) and each larger interval within the same spot containing candidate *attL*/*attR* (subject) was screened with BLASTn^96^, with the options *-evalue 0.001 -strand plus*. Alignments with ≥80% nucleotide identity were retained, and redundant hits fully contained within longer alignments were discarded. Pairs of hits on the subject sequence (large interval) aligning to overlapping regions of the query (short interval) were interpreted as duplication of a sequence segment associated with integration. The overlapping region on the shortest interval was defined as the putative *attB* site. The corresponding regions on the larger interval were defined as *attL* and *attR* sites (Fig S9A). Importantly, this approach makes no assumptions about the genomic context, such as the presence or position of the tyrosine recombinase genes.

To search for additional *att*-like sequences in the case of tandem integration events in the same interval (>1 *attL*/*R* per interval), we extracted the previously identified *attL* and *attR* sequences, along with the intergenic genomic region they delimit. Each *attL* and *attR* sequence (>15 bp) was then used as a query in a second BLASTn search against the extracted region to detect additional *att*-like sequences. Due to the short length of the *att* sequences (Fig S9B), BLASTn was run with parameters optimized for short queries *(-task blastn-short -evalue 1e-3 -strand plus*). Only hits with ≥80% identity and an alignment length exceeding 15 bp were retained. In cases of overlapping alignments, only the best-scoring hit was kept to remove redundancy. Retained *att* sites were then ordered according to their genomic coordinates, and *att*-delimited Genomic Intervals (GIs) were defined as the sequences larger than 2 Kb between successive *att* sites. This assumes that integration events occur in tandem relying on the same *att* site^66,119^, which is not always the case since some IMEs and other hitchers have been shown to integrate within their helpers^120,121^.

Most attachment sites are either not experimentally validated or unknown. To provide further evidence of the validity of the *ab initio* identification of *att* sites, we compared the predicted *att* sites with those reported by the TIGER/Islander database^47^, restricting the database to the 469,044 Genomic Islands identified in the same six species included in our study. Using the original *attL*/*attR* sequences, 99% of sites were recovered, either matching perfectly an *att* site described in the TIGER database or overlapping the sequences flanking the *att* sites (Fig S9C). However, because some *att* sites are very short (average of 21bp in our identification, 9bp in the TIGER database, as short as 2bp), comparisons based solely on their core sequences can yield spurious matches. To address this, each *att* sequence was extended together with its 10 nucleotides upstream and downstream (hereafter “extended *att*”), resulting in longer lengths. The 97% of our extended *att* sites were present in the TIGER database (Fig S9C). All sequences of the *att* sites, as well as the extended *att* sequences are available in Table S1.

### Building communities based on the weighted Gene Repertoire Relatedness (wGRR)

To assess the genetic relatedness between MGEs, we calculated the wGRR, using GRIS version 2.0 (https://gitlab.pasteur.fr/jgugliel/wgrr) with the default parameters as previously described^46^. Briefly, the software identifies the sequence similarities between all the proteins encoded in the elements using MMseqs2, version 16-747c6^104^, and retrieves the best bi-directional hits (BBH) between them. Once the BBHs have been defined, the wGRR can be calculated with the following formula,

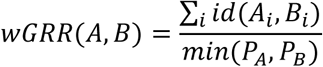

where *i* is the number of BBH between the two elements, *id(A_i_,B_i_)* is the identity of the BBH, and *P_A_* and *P_B_* are the number of proteins of the elements A and B, respectively. Hence, a wGRR=1 means that the two elements are identical (or one is entirely contained within the other), whereas wGRR=0 means the elements have no homologous protein-coding genes. To build genetically related clusters, we retrieved pairs of elements with wGRR>0.75 and built communities using the Louvain algorithm with the R package igraph, version 1.3.0 (https://CRAN.R-project.org/package=igraph)^122^, and the default resolution parameter (N=1). Graphs were visualized using the software Gephi, version 0.10.0^123^.

## Data availability

The dataset utilized for the current study is publicly available (RefSeq database, https://ftp.ncbi.nlm.nih.gov/genomes/refseq/, accessed in May 2023 and June 2025). The datasets generated are included in this published article and its supplementary information files. HMM profiles of the origins of transfer can be found at figshare^124^.

## Code availability

Custom R scripts to identify candidate *oriT*-containing regions for this study are available in figshare^125^.

## Acknowledgements

We thank all the Microbial Evolutionary Genomics Unit members for the regular discussions throughout the project. We also thank F. de la Cruz and M. P. Garcillán Barcia (IBBTEC-Universidad de Cantabria, Spain) for help and advice on this project. This work was funded by the INCEPTION project (PIA/ANR-16-CONV-0005, E.P.C.R.), the Fondation pour la Recherche Médicale (Equipe FRM, EQU201903007835, E.P.C.R.), the ANR RESIST (ANR 20 CE35 0014-02, E.P.C.R), the ANR TransfoConflict (ANR 20 CE12 0004 02), and the ANR LBX-62 IBEID and the ANR EcoRecEp (ANR-23-CE35-0006, M.T.). M.A.A was supported by a Pasteur-Roux-Cantarini fellowship of Institut Pasteur.

This work used the computational and storage services (TARS cluster) provided by the IT department at Institut Pasteur, Paris.

## References

1. Arnold, B. J., Huang, I.-T. & Hanage, W. P. Horizontal gene transfer and adaptive evolution in bacteria. Nat Rev Microbiol 20, 206–218 (2022).

2. De La Cruz, F., Frost, L. S., Meyer, R. J. & Zechner, E. L. Conjugative DNA metabolism in Gram-negative bacteria. FEMS Microbiol Rev 34, 18–40 (2010).

3. Virolle, C., Goldlust, K., Djermoun, S., Bigot, S. & Lesterlin, C. Plasmid Transfer by Conjugation in Gram-Negative Bacteria: From the Cellular to the Community Level. Genes 11, 1239 (2020).

4. Smillie, C., Garcillán-Barcia, M. P., Francia, M. V., Rocha, E. P. C. & de la Cruz, F. Mobility of Plasmids. Microbiol Mol Biol Rev 74, 434–452 (2010).

5. Waldor, M. K. Mobilizable genomic islands: going mobile with *oriT* mimicry. Molecular Microbiology 78, 537–540 (2010).

6. Ares-Arroyo, M., Coluzzi, C. & Rocha, E. P. C. Origins of transfer establish networks of functional dependencies for plasmid transfer by conjugation. Nucleic Acids Research 51, 3001–3016 (2023).

7. Lang, A. S., Buchan, A. & Burrus, V. Interactions and evolutionary relationships among bacterial mobile genetic elements. Nat Rev Microbiol 23, 423–438 (2025).

8. Ares-Arroyo, M., Coluzzi, C., Moura De Sousa, J. A. & Rocha, E. P. C. Hijackers, hitchhikers, or co-drivers? The mysteries of mobilizable genetic elements. PLoS Biol 22, e3002796 (2024).

9. Burrus, V. & Waldor, M. K. Shaping bacterial genomes with integrative and conjugative elements. Research in Microbiology 155, 376–386 (2004).

10. Johnson, C. M. & Grossman, A. D. Integrative and Conjugative Elements (ICEs): What They Do and How They Work. Annu. Rev. Genet. 49, 577–601 (2015).

11. Audrey, B., Cellier, N., White, F., Jacques, P.-É. & Burrus, V. A systematic approach to classify and characterize genomic islands driven by conjugative mobility using protein signatures. Nucleic Acids Research 51, 8402–8412 (2023).

12. Brochet, M. et al. Atypical association of DDE transposition with conjugation specifies a new family of mobile elements. Molecular Microbiology 71, 948–959 (2009).

13. Guglielmini, J., de la Cruz, F. & Rocha, E. P. C. Evolution of Conjugation and Type IV Secretion Systems. Molecular Biology and Evolution 30, 315–331 (2013).

14. Guglielmini, J., Quintais, L., Garcillán-Barcia, M. P., De La Cruz, F. & Rocha, E. P. C. The Repertoire of ICE in Prokaryotes Underscores the Unity, Diversity, and Ubiquity of Conjugation. PLoS Genet 7, e1002222 (2011).

15. Guédon, G., Libante, V., Coluzzi, C., Payot, S. & Leblond-Bourget, N. The Obscure World of Integrative and Mobilizable Elements, Highly Widespread Elements that Pirate Bacterial Conjugative Systems. Genes 8, 337 (2017).

16. Botelho, J. Defense systems are pervasive across chromosomally integrated mobile genetic elements and are inversely correlated to virulence and antimicrobial resistance. Nucleic Acids Research 51, 4385–4397 (2023).

17. Ramsay, J. P. & Firth, N. Diverse mobilization strategies facilitate transfer of non-conjugative mobile genetic elements. Current Opinion in Microbiology 38, 1–9 (2017).

18. Achard, A. & Leclercq, R. Characterization of a Small Mobilizable Transposon, MTnSag1, in *Streptococcus agalactiae*. J Bacteriol 189, 4328–4331 (2007).

19. Carraro, N., Rivard, N., Ceccarelli, D., Colwell, R. R. & Burrus, V. IncA/C Conjugative Plasmids Mobilize a New Family of Multidrug Resistance Islands in Clinical Vibrio cholerae Non-O1/Non-O139 Isolates from Haiti. mBio 7, e00509–16 (2016).

20. Daccord, A., Ceccarelli, D., Rodrigue, S. & Burrus, V. Comparative Analysis of Mobilizable Genomic Islands. Journal of Bacteriology 195, 606–614 (2013).

21. Botelho, J. & Schulenburg, H. The Role of Integrative and Conjugative Elements in Antibiotic Resistance Evolution. Trends in Microbiology 29, 8–18 (2021).

22. Colombi, E. et al. Comparative analysis of integrative and conjugative mobile genetic elements in the genus Mesorhizobium. Microbial Genomics 7, (2021).

23. LeGault, K. N. et al. Temporal shifts in antibiotic resistance elements govern phage-pathogen conflicts. Science 373, eabg2166 (2021).

24. Seth-Smith, H. M. B. et al. Structure, Diversity, and Mobility of the Salmonella Pathogenicity Island 7 Family of Integrative and Conjugative Elements within Enterobacteriaceae. J Bacteriol 194, 1494–1504 (2012).

25. Jones, J. M., Grinberg, I., Eldar, A. & Grossman, A. D. A mobile genetic element increases bacterial host fitness by manipulating development. eLife 10, e65924 (2021).

26. Bean, E. L., McLellan, L. K. & Grossman, A. D. Activation of the integrative and conjugative element Tn916 causes growth arrest and death of host bacteria. PLoS Genet 18, e1010467 (2022).

27. Reinhard, F., Miyazaki, R., Pradervand, N. & van der Meer, J. R. Cell Differentiation to “Mating Bodies” Induced by an Integrating and Conjugative Element in Free-Living Bacteria. Current Biology 23, 255–259 (2013).

28. Boyd, E. F., Almagro-Moreno, S. & Parent, M. A. Genomic islands are dynamic, ancient integrative elements in bacterial evolution. Trends in Microbiology 17, 47–53 (2009).

29. Juhas, M. et al. Genomic islands: tools of bacterial horizontal gene transfer and evolution. FEMS Microbiol Rev 33, 376–393 (2009).

30. Hacker, J. & Kaper, J. B. Pathogenicity Islands and the Evolution of Microbes. Annu. Rev. Microbiol. 54, 641–679 (2000).

31. Sia, C. M., Pearson, J. S., Howden, B. P., Williamson, D. A. & Ingle, D. J. Salmonella pathogenicity islands in the genomic era. Trends in Microbiology 33, 752–764 (2025).

32. Doublet, B., Boyd, D., Mulvey, M. R. & Cloeckaert, A. The *Salmonella* genomic island 1 is an integrative mobilizable element. Molecular Microbiology 55, 1911–1924 (2005).

33. Durrant, M. G., Li, M. M., Siranosian, B. A., Montgomery, S. B. & Bhatt, A. S. A Bioinformatic Analysis of Integrative Mobile Genetic Elements Highlights Their Role in Bacterial Adaptation. Cell Host & Microbe 27, 140–153.e9 (2020).

34. Hussain, F. A. et al. Rapid evolutionary turnover of mobile genetic elements drives bacterial resistance to phages. Science 374, 488–492 (2021).

35. Ares-Arroyo, M., Nucci, A. & Rocha, E. P. C. Expanding the diversity of origin of transfer-containing sequences in mobilizable plasmids. Nat Microbiol 9, 3240–3253 (2024).

36. Guglielmini, J. et al. Key components of the eight classes of type IV secretion systems involved in bacterial conjugation or protein secretion. Nucleic Acids Research 42, 5715–5727 (2014).

37. Kiss, J. et al. Identification and Characterization of oriT and Two Mobilization Genes Required for Conjugative Transfer of Salmonella Genomic Island 1. Front. Microbiol. 10, 457 (2019).

38. Lorenzo-Díaz, F., Fernández-López, C., Guillén-Guío, B., Bravo, A. & Espinosa, M. Relaxase MobM Induces a Molecular Switch at Its Cognate Origin of Transfer. Front. Mol. Biosci. 5, 17 (2018).

39. Nowak, K. P. et al. Molecular and Functional Characterization of MobK Protein—A Novel-Type Relaxase Involved in Mobilization for Conjugational Transfer of Klebsiella pneumoniae Plasmid pIGRK. IJMS 22, 5152 (2021).

40. Wisniewski, J. A. et al. TcpM: a novel relaxase that mediates transfer of large conjugative plasmids from *Clostridium perfringens*. Molecular Microbiology 99, 884–896 (2016).

41. Oliveira, P. H., Touchon, M., Cury, J. & Rocha, E. P. C. The chromosomal organization of horizontal gene transfer in bacteria. Nat Commun 8, 841 (2017).

42. Garcillán-Barcia, M. P., de la Cruz, F. & Rocha, E. P. C. The extended mobility of plasmids. Nucleic Acids Research 53, gkaf652 (2025).

43. Fekete, R. A. & Frost, L. S. Mobilization of Chimeric *oriT* Plasmids by F and R100-1: Role of Relaxosome Formation in Defining Plasmid Specificity. J Bacteriol 182, 4022–4027 (2000).

44. Avila, P., Núñez, B. & de la Cruz, F. Plasmid R6K contains two functional oriTs which can assemble simultaneously in relaxosomes in vivo. J Mol Biol 261, 135–143 (1996).

45. Miyazaki, R. & Van Der Meer, J. R. A dual functional origin of transfer in the ICEclc genomic island of Pseudomonas knackmussii B13: oriT of ICEclc. Molecular Microbiology 79, 743–758 (2011).

46. Cury, J., Oliveira, P. H., de la Cruz, F. & Rocha, E. P. C. Host Range and Genetic Plasticity Explain the Coexistence of Integrative and Extrachromosomal Mobile Genetic Elements. Molecular Biology and Evolution 35, 2230–2239 (2018).

47. Yu, S. L. et al. The TIGER/Islander resource for genomic islands. Microbiol Resour Announc 14, e00492–25 (2025).

48. Cury, J., Touchon, M. & Rocha, E. P. C. Integrative and conjugative elements and their hosts: composition, distribution and organization. Nucleic Acids Res 45, 8943–8956 (2017).

49. Coluzzi, C., Garcillán-Barcia, M. P., de la Cruz, F. & Rocha, E. P. C. Evolution of Plasmid Mobility: Origin and Fate of Conjugative and Nonconjugative Plasmids. Molecular Biology and Evolution 39, msac115 (2022).

50. Daccord, A., Ceccarelli, D. & Burrus, V. Integrating conjugative elements of the SXT/R391 family trigger the excision and drive the mobilization of a new class of Vibrio genomic islands: ICE-mediated GI mobilization. Molecular Microbiology 78, 576–588 (2010).

51. Douard, G., Praud, K., Cloeckaert, A. & Doublet, B. The Salmonella Genomic Island 1 Is Specifically Mobilized In Trans by the IncA/C Multidrug Resistance Plasmid Family. PLoS ONE 5, e15302 (2010).

52. Geng, P. et al. Horizontal transfer of large plasmid with type IV secretion system and mosquitocidal genomic island with excision and integration capabilities in *Lysinibacillus sphaericus*. Environmental Microbiology 23, 5131–5146 (2021).

53. Valentine, P. J., Shoemaker, N. B. & Salyers, A. A. Mobilization of Bacteroides plasmids by Bacteroides conjugal elements. J Bacteriol 170, 1319–1324 (1988).

54. Wong, J. et al. The assembly of a hybrid type IV secretion system by a Crohn’s disease-associated Escherichia coli strain. Nat Commun 16, 8797 (2025).

55. Taylor, D. E. et al. Genomic Variability of O Islands Encoding Tellurite Resistance in Enterohemorrhagic *Escherichia coli* O157:H7 Isolates. J Bacteriol 184, 4690–4698 (2002).

56. Neupane, D. P., Bearson, B. L. & Bearson, S. M. D. Localization of the origin of transfer (oriT) for Salmonella Genomic Island 4 (SGI-4) from Salmonella enterica serovar I 4, [5],12:i:-. DNA Research dsaf023 (2025) doi:10.1093/dnares/dsaf023.

57. Miruna Perta, A., et al. Integrative mobilizable elements are pervasive throughout Pseudomonadota. Preprint at 10.64898/2026.01.12.698999 (2026).

58. Lin, T.-L., Lee, C.-Z., Hsieh, P.-F., Tsai, S.-F. & Wang, J.-T. Characterization of Integrative and Conjugative Element ICE *Kp1*-Associated Genomic Heterogeneity in a *Klebsiella pneumoniae* Strain Isolated from a Primary Liver Abscess. J Bacteriol 190, 515–526 (2008).

59. Beamud, B., Benz, F. & Bikard, D. Going viral: The role of mobile genetic elements in bacterial immunity. Cell Host & Microbe 32, 804–819 (2024).

60. Shaw, L. P., Rocha, E. P. C. & MacLean, R. C. Restriction-modification systems have shaped the evolution and distribution of plasmids across bacteria. Nucleic Acids Research 51, 6806–6818 (2023).

61. Jaskólska, M., Adams, D. W. & Blokesch, M. Two defence systems eliminate plasmids from seventh pandemic Vibrio cholerae. Nature 604, 323–329 (2022).

62. Couturier, A. et al. Real-time visualisation of the intracellular dynamics of conjugative plasmid transfer. Nat Commun 14, 294 (2023).

63. Samuel, B., Mittelman, K., Croitoru, S. Y., Ben Haim, M. & Burstein, D. Diverse anti-defence systems are encoded in the leading region of plasmids. Nature 635, 186–192 (2024).

64. Marcoleta, A. E., Berríos-Pastén, C., Nuñez, G., Monasterio, O. & Lagos, R. Klebsiella pneumoniae Asparagine tDNAs Are Integration Hotspots for Different Genomic Islands Encoding Microcin E492 Production Determinants and Other Putative Virulence Factors Present in Hypervirulent Strains. Front. Microbiol. 7, (2016).

65. Rivard, N., Colwell, R. R. & Burrus, V. Antibiotic Resistance in Vibrio cholerae: Mechanistic Insights from IncC Plasmid-Mediated Dissemination of a Novel Family of Genomic Islands Inserted at *trmE*. mSphere 5, 10.1128/msphere.00748-20 (2020).

66. Bellanger, X. et al. Site-specific accretion of an integrative conjugative element together with a related genomic island leads to *cis* mobilization and gene capture. Molecular Microbiology 81, 912–925 (2011).

67. Ray, M. D., Boundy, S. & Archer, G. L. Transfer of the methicillin resistance genomic island among staphylococci by conjugation. Molecular Microbiology 100, 675–685 (2016).

68. Libante, V. et al. Mobilization of IMEs Integrated in the oriT of ICEs Involves Their Own Relaxase Belonging to the Rep-Trans Family of Proteins. Genes 11, 1004 (2020).

69. Hua, M. et al. A chromosome-encoded T4SS independently contributes to horizontal gene transfer in Enterococcus faecalis. Cell Reports 41, 111609 (2022).

70. Whittle, G., Hamburger, N., Shoemaker, N. B. & Salyers, A. A. A Bacteroides Conjugative Transposon, CTnERL, Can Transfer a Portion of Itself by Conjugation without Excising from the Chromosome. J Bacteriol 188, 1169–1174 (2006).

71. Deep, A. et al. The SMC-family Wadjet complex protects bacteria from plasmid transformation by recognition and cleavage of closed-circular DNA. Molecular Cell 82, 4145–4159.e7 (2022).

72. Zongo, P. D. et al. An antiplasmid system drives antibiotic resistance gene integration in carbapenemase-producing Escherichia coli lineages. Nat Commun 15, 4093 (2024).

73. Redondo-Salvo, S. et al. Pathways for horizontal gene transfer in bacteria revealed by a global map of their plasmids. Nat Commun 11, 3602 (2020).

74. Carraro, N., Poulin, D. & Burrus, V. Replication and Active Partition of Integrative and Conjugative Elements (ICEs) of the SXT/R391 Family: The Line between ICEs and Conjugative Plasmids Is Getting Thinner. PLoS Genet 11, e1005298 (2015).

75. Delavat, F., Miyazaki, R., Carraro, N., Pradervand, N. & Van Der Meer, J. R. The hidden life of integrative and conjugative elements. FEMS Microbiology Reviews 41, 512–537 (2017).

76. Meyer, R. The r1162 mob proteins can promote conjugative transfer from cryptic origins in the bacterial chromosome. J Bacteriol 191, 1574–1580 (2009).

77. Ramsay, J. P. et al. A widely conserved molecular switch controls quorum sensing and symbiosis island transfer in *M esorhizobium loti* through expression of a novel antiactivator. Molecular Microbiology 87, 1–13 (2013).

78. Burrus, V. & Waldor, M. K. Control of SXT Integration and Excision. J Bacteriol 185, 5045–5054 (2003).

79. Waters, J. L. & Salyers, A. A. Regulation of CTnDOT Conjugative Transfer Is a Complex and Highly Coordinated Series of Events. mBio 4, e00569–13 (2013).

80. Verdonk, C. J. et al. Structural basis for control of integrative and conjugative element excision and transfer by the oligomeric winged helix–turn–helix protein RdfS. Nucleic Acids Research 53, gkaf249 (2025).

81. Pfeifer, E. & Rocha, E. P. C. Phage-plasmids promote recombination and emergence of phages and plasmids. Nat Commun 15, 1545 (2024).

82. Wozniak, R. A. F. & Waldor, M. K. Integrative and conjugative elements: mosaic mobile genetic elements enabling dynamic lateral gene flow. Nat Rev Microbiol 8, 552–563 (2010).

83. Jacob, F. & Wollman, E. L. Genetic and physical determinations of chromosomal segments in Escherichia coli. Symp Soc Exp Biol 12, 75–92 (1958).

84. Errington, J. & Pughe, N. Upper limit for DNA packaging by Bacillus subtilis bacteriophage ϕ 105: Isolation of phage deletion mutants by induction of oversized prophages. Molec Gen Genet 210, 347–351 (1987).

85. Humphrey, S. et al. Staphylococcal phages and pathogenicity islands drive plasmid evolution. Nat Commun 12, 5845 (2021).

86. Coluzzi, C. & Rocha, E. P. C. The Spread of Antibiotic Resistance Is Driven by Plasmids Among the Fastest Evolving and of Broadest Host Range. Molecular Biology and Evolution 42, msaf060 (2025).

87. Zheng, Q., Li, L., Yin, X., Che, Y. & Zhang, T. Is ICE hot? A genomic comparative study reveals integrative and conjugative elements as “hot” vectors for the dissemination of antibiotic resistance genes. mSystems 8, e00178–23 (2023).

88. Enault, F. et al. Phages rarely encode antibiotic resistance genes: a cautionary tale for virome analyses. The ISME Journal 11, 237–247 (2017).

89. Pfeifer, E., Bonnin, R. & Rocha, E. P. C. Phage-Plasmids Spread Antibiotic Resistance Genes through Infection and Lysogenic Conversion. http://biorxiv.org/lookup/doi/10.1101/2022.06.24.497495 (2022) doi:10.1101/2022.06.24.497495.

90. Hyatt, D. et al. Prodigal: prokaryotic gene recognition and translation initiation site identification. BMC Bioinformatics 11, 119 (2010).

91. Cantalapiedra, C. P., Hernández-Plaza, A., Letunic, I., Bork, P. & Huerta-Cepas, J. eggNOG-mapper v2: Functional Annotation, Orthology Assignments, and Domain Prediction at the Metagenomic Scale. Molecular Biology and Evolution 38, 5825–5829 (2021).

92. Néron, B., et al. MacSyFinder v2: Improved modelling and search engine to identify molecular systems in genomes. Peer Community Journal 3, e28 (2023).

93. Tesson, F. et al. Systematic and quantitative view of the antiviral arsenal of prokaryotes. Nat Commun 13, 2561 (2022).

94. Tesson, F. et al. A Comprehensive Resource for Exploring Antiphage Defense: DefenseFinder Webservice, Wiki and Databases. Peer Community Journal 4, e91 (2024).

95. Feldgarden, M. et al. AMRFinderPlus and the Reference Gene Catalog facilitate examination of the genomic links among antimicrobial resistance, stress response, and virulence. Sci Rep 11, 12728 (2021).

96. Camacho, C. et al. BLAST+: architecture and applications. BMC Bioinformatics 10, 421 (2009).

97. Zhou, S., Liu, B., Zheng, D., Chen, L. & Yang, J. VFDB 2025: an integrated resource for exploring anti-virulence compounds. Nucleic Acids Research 53, D871–D877 (2025).

98. Xie, Z. & Tang, H. ISEScan: automated identification of insertion sequence elements in prokaryotic genomes. Bioinformatics 33, 3340–3347 (2017).

99. Mistry, J. et al. Pfam: The protein families database in 2021. Nucleic Acids Res 49, D412–D419 (2021).

100. Liu, G. et al. oriTDB: a database of the origin-of-transfer regions of bacterial mobile genetic elements. Nucleic Acids Research 53, D163–D168 (2025).

101. Núñez, B. & De La Cruz, F. Two atypical mobilization proteins are involved in plasmid CloDF13 relaxation. Molecular Microbiology 39, 1088–1099 (2001).

102. Francia, M. V. & Clewell, D. B. Transfer origins in the conjugative *Enterococcus faecalis* plasmids pAD1 and pAM373: identification of the pAD1 *nic* site, a specific relaxase and a possible TraG-like protein. Molecular Microbiology 45, 375–395 (2002).

103. Pearson, W. R. & Lipman, D. J. Improved tools for biological sequence comparison. Proc. Natl. Acad. Sci. U.S.A. 85, 2444–2448 (1988).

104. Steinegger, M. & Söding, J. MMseqs2 enables sensitive protein sequence searching for the analysis of massive data sets. Nat Biotechnol 35, 1026–1028 (2017).

105. Katoh, K. & Standley, D. M. MAFFT Multiple Sequence Alignment Software Version 7: Improvements in Performance and Usability. Molecular Biology and Evolution 30, 772–780 (2013).

106. Carraro, N. et al. Development of pVCR94ΔX from Vibrio cholerae, a prototype for studying multidrug resistant IncA/C conjugative plasmids. Front. Microbiol. 5, 5:44 (2014).

107. Ceccarelli, D., Daccord, A., René, M. & Burrus, V. Identification of the Origin of Transfer (*oriT*) and a New Gene Required for Mobilization of the SXT/R391 Family of Integrating Conjugative Elements. J Bacteriol 190, 5328–5338 (2008).

108. Tian, D. et al. Prevalence of hypervirulent and carbapenem-resistant *Klebsiella pneumoniae* under divergent evolutionary patterns. Emerging Microbes & Infections 11, 1936–1949 (2022).

109. Shen, W., Le, S., Li, Y. & Hu, F. SeqKit: A Cross-Platform and Ultrafast Toolkit for FASTA/Q File Manipulation. PLoS ONE 11, e0163962 (2016).

110. Wheeler, T. J. & Eddy, S. R. nhmmer: DNA homology search with profile HMMs. Bioinformatics 29, 2487–2489 (2013).

111. Fricke, W. F. et al. Comparative Genomics of the IncA/C Multidrug Resistance Plasmid Family. J Bacteriol 191, 4750–4757 (2009).

112. Hegyi, A., Szabó, M., Olasz, F. & Kiss, J. Identification of oriT and a recombination hot spot in the IncA/C plasmid backbone. Sci Rep 7, 10595 (2017).

113. Ilchenko, K., Bonnin, R. A., Rocha, E. P. & Pfeifer, E. Efficient detection and typing of phage-plasmids. Preprint at 10.1101/2025.08.29.673033 (2025).

114. Redondo-Salvo, S. et al. COPLA, a taxonomic classifier of plasmids. BMC Bioinformatics 22, 390 (2021).

115. Garcillán-Barcia, M. P., Redondo-Salvo, S. & De La Cruz, F. Plasmid classifications. Plasmid 126, 102684 (2023).

116. Garcillán-Barcia, M. P., Francia, M. V. & de La Cruz, F. The diversity of conjugative relaxases and its application in plasmid classification. FEMS Microbiol Rev 33, 657–687 (2009).

117. Perrin, A. & Rocha, E. P. C. PanACoTA: a modular tool for massive microbial comparative genomics. NAR Genom Bioinform 3, lqaa106 (2021).

118. Rajeev, L., Malanowska, K. & Gardner, J. F. Challenging a Paradigm: the Role of DNA Homology in Tyrosine Recombinase Reactions. Microbiol Mol Biol Rev 73, 300–309 (2009).

119. Pavlovic, G., Burrus, V., Gintz, B., Decaris, B. & Guédon, G. Evolution of genomic islands by deletion and tandem accretion by site-specific recombination: ICESt1-related elements from Streptococcus thermophilus. Microbiology 150, 759–774 (2004).

120. Ambroset, C. et al. New Insights into the Classification and Integration Specificity of Streptococcus Integrative Conjugative Elements through Extensive Genome Exploration. Front. Microbiol. 6, (2016).

121. Tommasini, D., Mageeney, C. M. & Williams, K. P. Helper-embedded satellites from an integrase clade that repeatedly targets prophage late genes. NAR Genomics and Bioinformatics 5, lqad036 (2023).

122. Csárdi, G., et al. igraph for R: R interface of the igraph library for graph theory and network analysis. Zenodo 10.5281/ZENODO.7682609 (2025).

123. Bastian, M., Heymann, S. & Jacomy, M. Gephi: An Open Source Software for Exploring and Manipulating Networks. ICWSM 3, 361–362 (2009).

124. Ares-Arroyo, M. HMM profiles for the screening of conjugative origins of transfer. 8143879 Bytes figshare 10.6084/M9.FIGSHARE.31047685 (2026).

125. Ares-Arroyo, M. R Scripts for the identification of sequences containing origins of transfer by conjugation. 37685 Bytes figshare 10.6084/M9.FIGSHARE.26870155.V1 (2024).

