## Supplementary Figures for "Genomic Islands as minimal hitchers of conjugative elements" (from another version of this preprint: version 1, 2026-01-13)

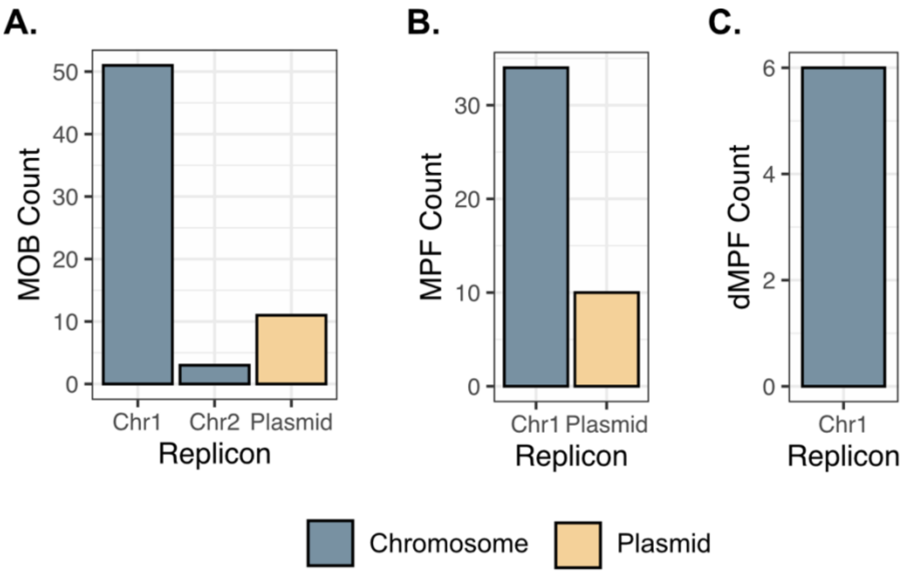


**Figure S1. A.** Relaxases identified across the replicons of *V. cholerae*. Chr1: First chromosome. Chr2. Second chromosome. **B.** Complete Mating Pair Formation (MPF) systems identified. **C.** Decayed incomplete MPF systems.

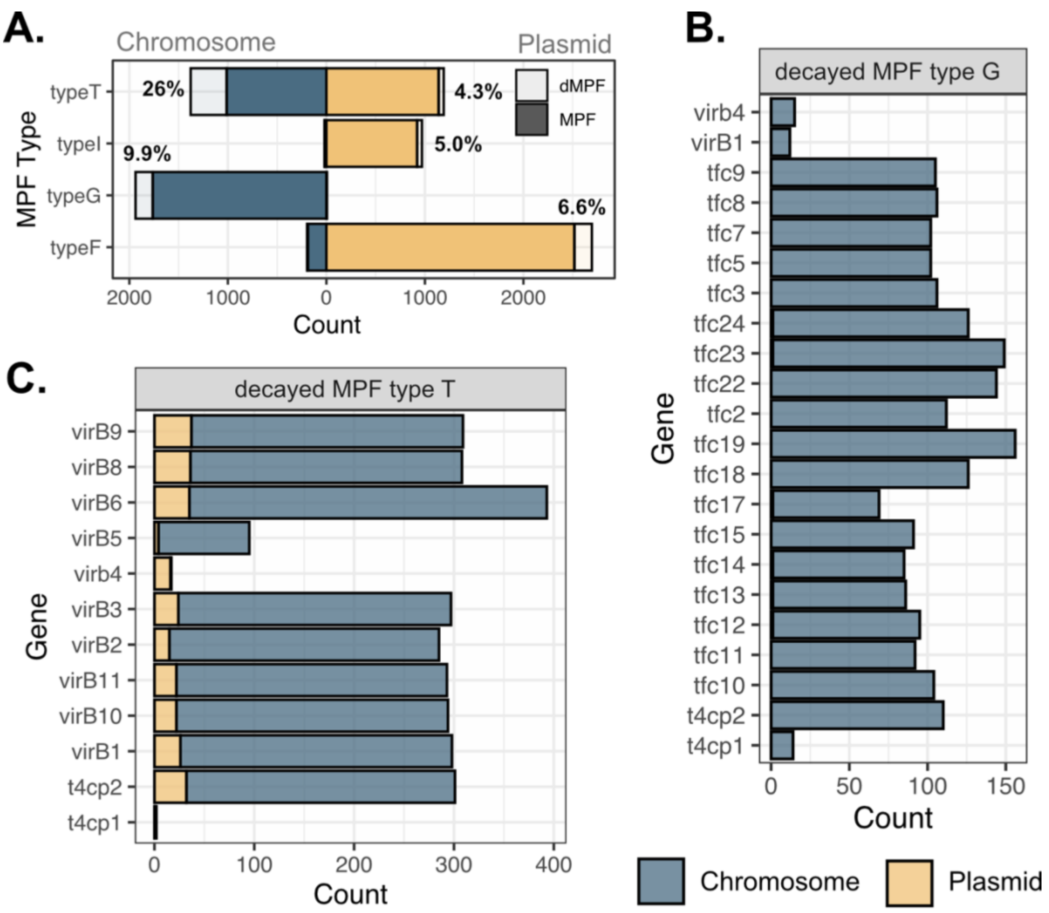


**Figure S2. A.** MPF types identified in chromosomes (left) and plasmids (right). The clearest area, and the percentage next to it, indicates the fraction of decayed MPF systems per MPF type and replicon. **B.** and **C.** Protein coding genes present in the decayed MPF systems of type G (**B.**) and type T (**C.**). The color indicates the replicon where the system is encoded: blue in chromosome, orange in plasmids.

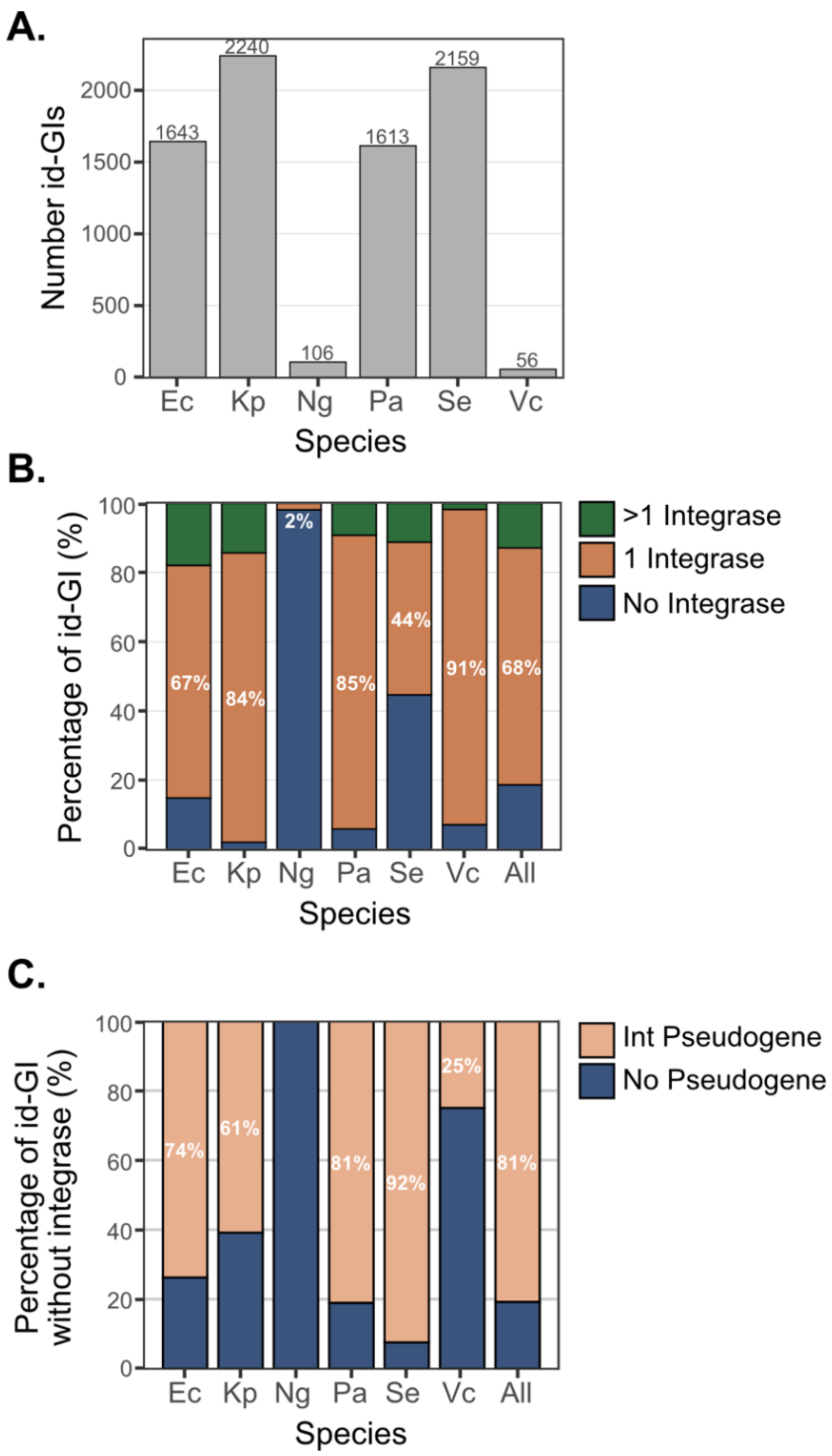


**Figure S3. A.** Number of interval-delimited Genomic Islands identified per species. **B.** Percentage of interval-delimited Genomic Islands harboring 0 integrase genes, 1 integrase genes, or >1 integrase genes. The text within the bar indicates the percentage of id-GIs with 1 integrase. **C.** Percentage of id-GI without integrases, where a integrase pseudogene was identified.

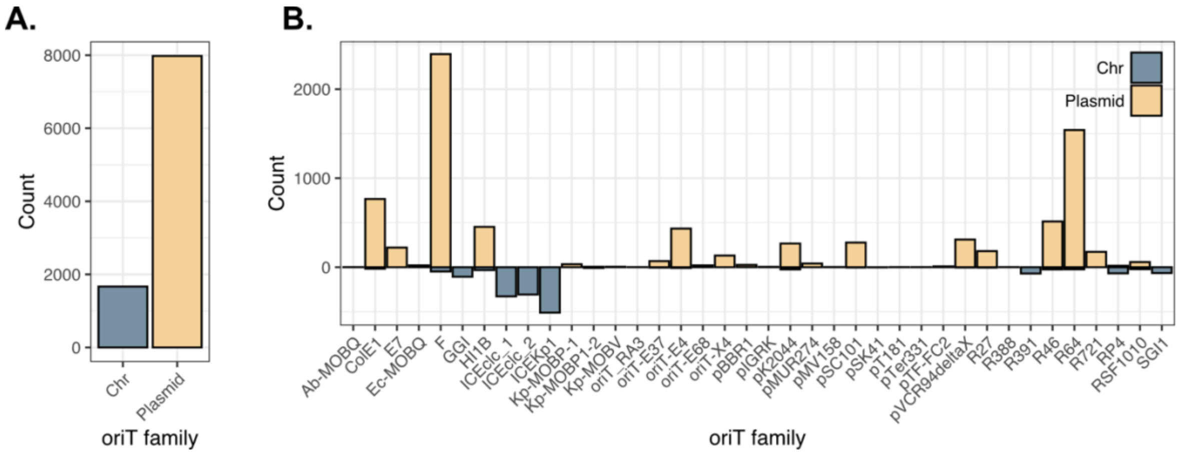


**Figure S4. A.** Number of known *oriTs* identified in chromosomes (Chr) and plasmids. **B.** Number of each *oriT* family identified in plasmids (light grey, up) and chromosomes (dark grey, bottom).

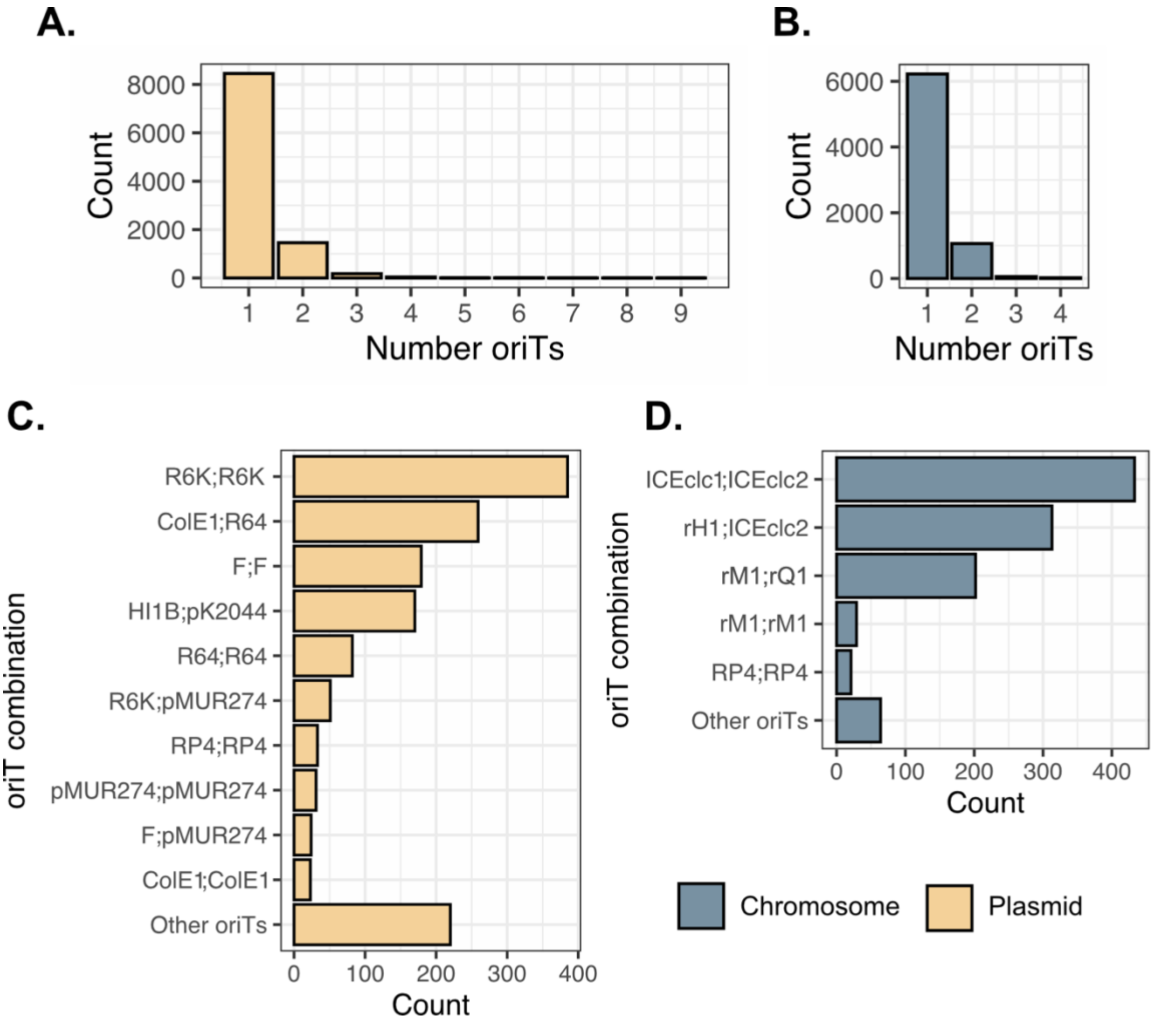


**Figure S5. A.** Distribution of *oriTs* per plasmid. **B.** Distribution of *oriTs* per non-*att*-delimited Genomic Island. **C.** *oriT* combinations in plasmids carrying 2 *oriTs*. **D.** *oriT* combinations in non-delimited Genomic Islands carrying 2 *oriTs*.

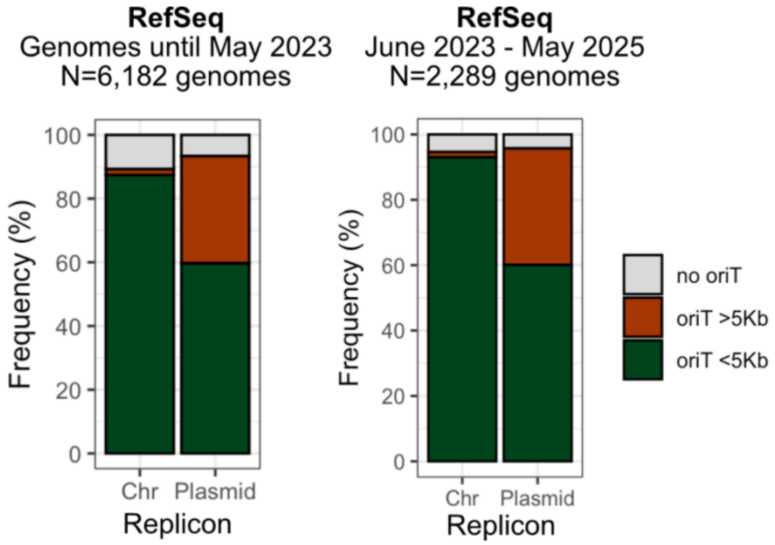


**Figure S6.** Fraction of relaxases with an *oriT* in the surrounding 5Kb (green), farther than 5Kb, or no *oriT* identified (grey) across the dataset used to inferred the *oriTs* (left, complete genomes from RefSeq before May 2023) and a new dataset (right, complete genomes from RefSeq between June 2022 and May 2025).

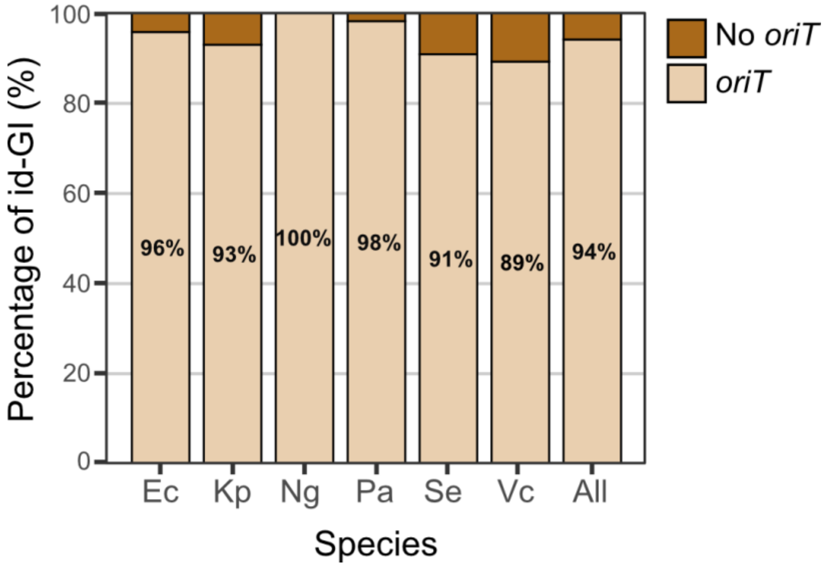


**Figure S7.** Percentage of interval-delimited Genomic Island with an origin of transfer.

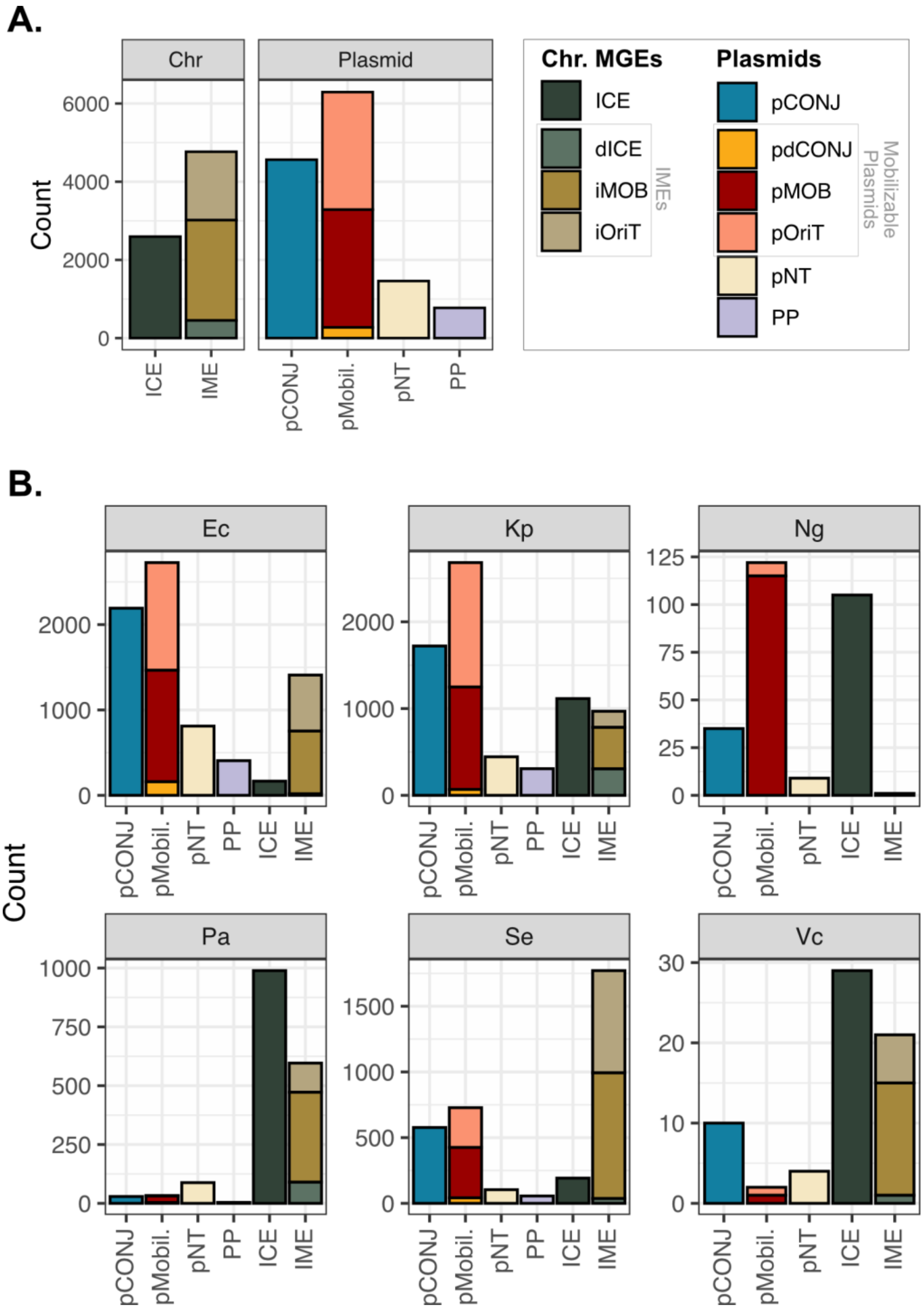


**Figure S8. A.** Number of MGEs across the six species in both chromosomes (Chr) and plasmids. The color of the elements is represented at the right of the figure. pNT: non-transmissible plasmid. PP: Phage-Plasmid. **B.** Number of each element per species. Ec: *E. coli*. Kp: *K. pneumoniae*. Ng. *N. gonhorroeae*. Pa. *P. aeruginosa*. Se. *S. enterica*. Vc. *V. cholerae*.

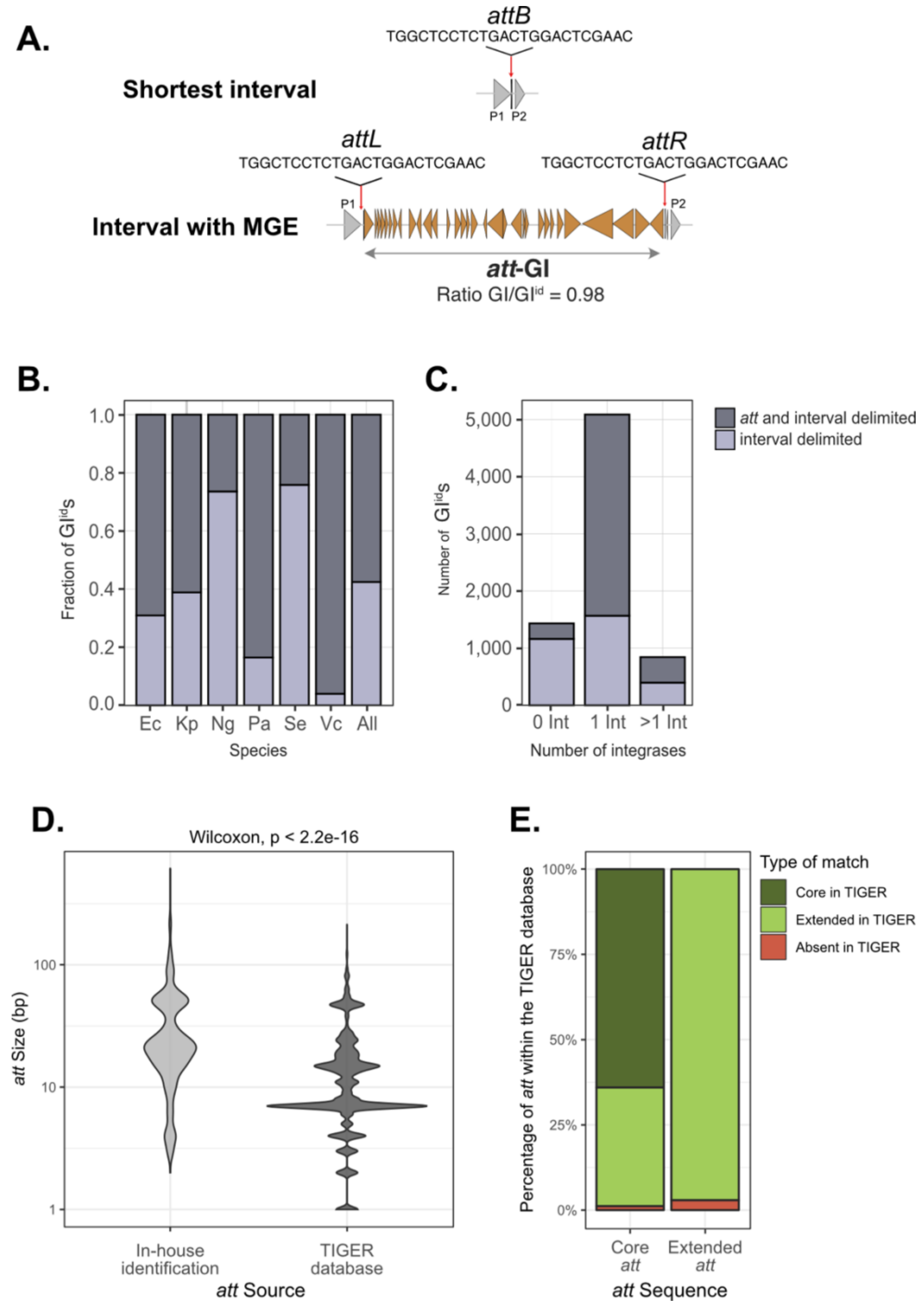


**Figure S9. A.** Definition of *attB, attL,* and *attR*. P1 and P2 are consecutive persistent gene families. **B.** Fraction of interval-delimited Genomic Islands where an *att*-delimited GI could be identified. **C.** Number of interval-delimited Genomic Islands with 0, 1 or >1 tyrosine recombinase. The color indicates if an *att*-GI was identified within the id-GI. **D.** Size of the *att* sites identified by our method and the ones present in the TIGER database. **E.** Percentage of *att* sites identified present in the TIGER database. Because some *att* sites are very short (>2bp), comparisons based solely on their "core *att*" sequences can yield spurious matches. Each *att* sequence was extended together with its 10 nucleotides upstream and downstream ("extended *att*"), resulting in longer lengths.

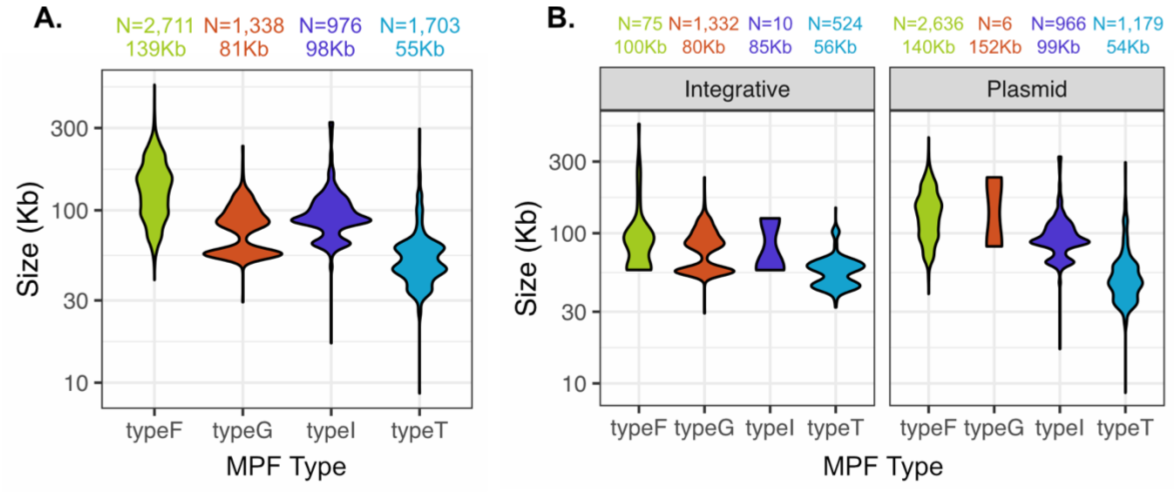


**Figure S10. A.** Size distribution of the MGE carrying a complete MPF system. Numbers on top indicate the number of elements, and the mean size. **B.** Size distribution split into ICEs (integrative, left) and conjugative plasmids (Plasmid, right).

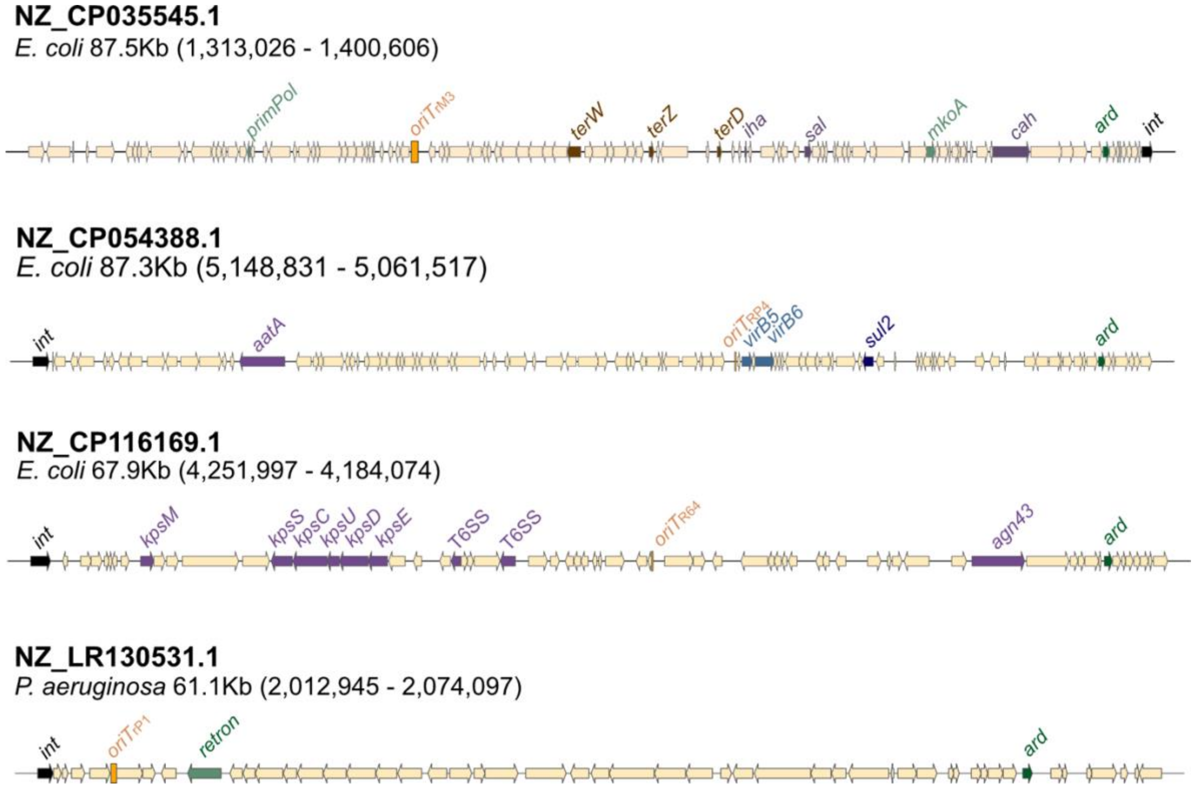


**Figure S11.** Genetic map of examples of large *att*-delimited iOriT elements. Genes in green represent defense systems, brown tellurite resistance, purple virulence, black integrase, and orange the *oriT*.

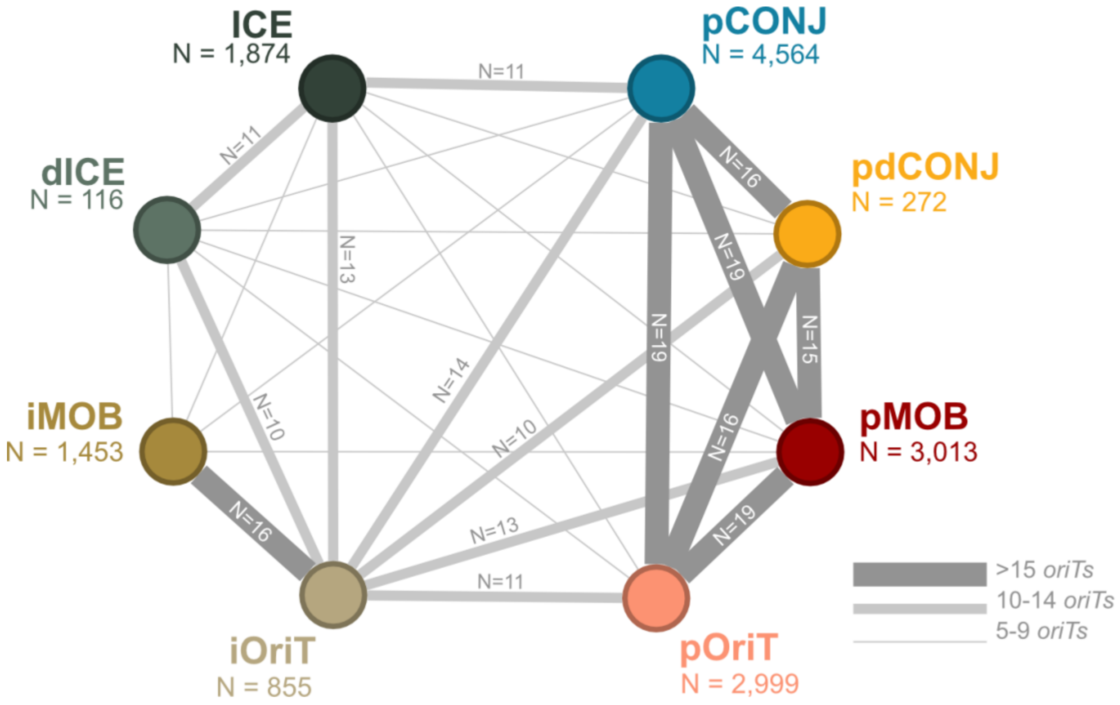


**Figure S12.** *oriT* families shared between different MGE types. The nodes of the graph represent the MGEs, their size correlated to the sample size. Integrative MGEs are shown to the left, episomal MGEs to the right. Edges connect MGE types when a same *oriT* has been found in both elements, its width representing the number of different *oriTs* in common. The legend is at the bottom right of the figure. When they share >5 *oriT* families, the number is shown next to the edge.

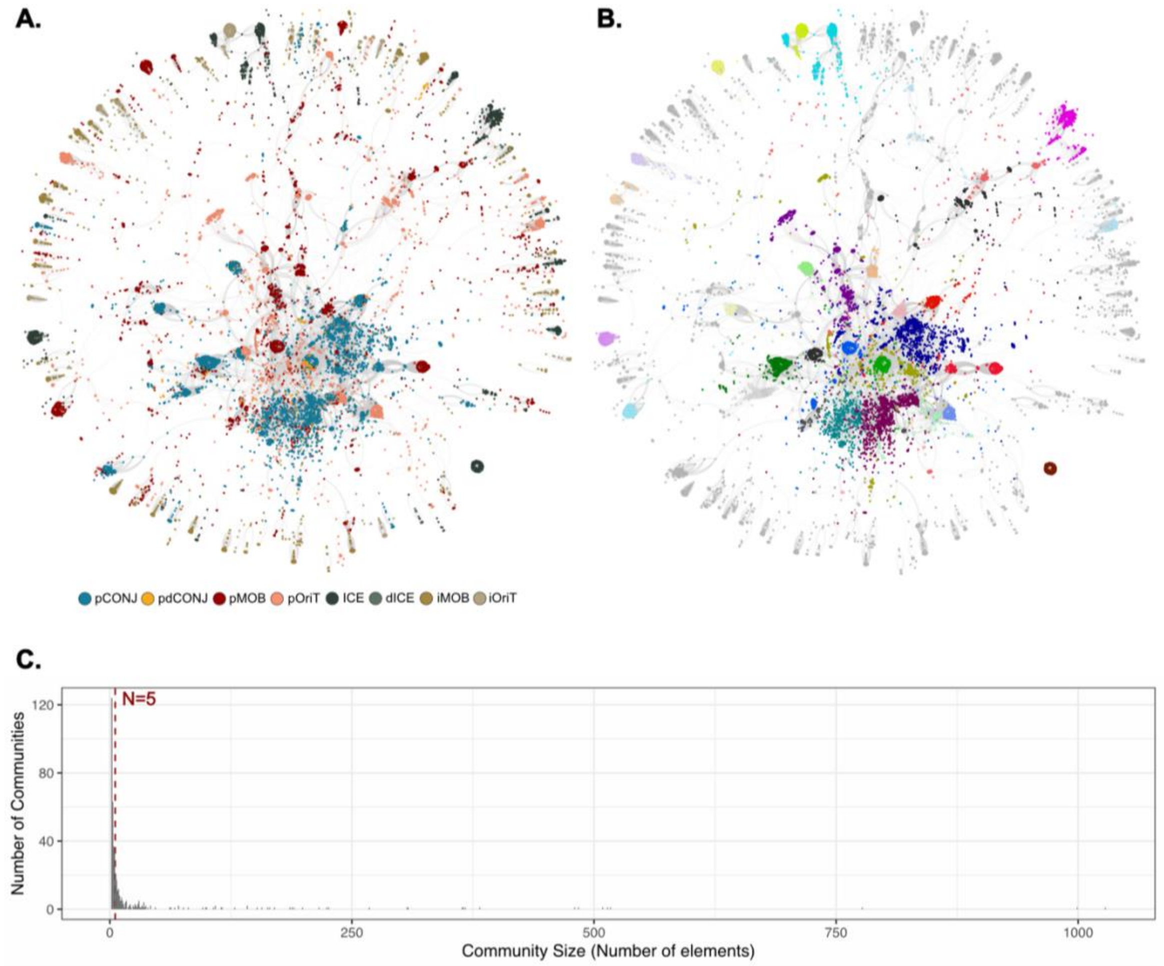


**Figure S13. A.** MGE network according to the wGRR. Each node is an MGE. Nodes are connected when the wGRR between the MGEs is >0.75. The color of the nodes represents the MGE type, being its legend at the bottom of the figure. **B.** Same as the graph A. but each node is colored according to the community it belongs. Only the major communities are colored. **C.** Size of the communities. The vertical red dashed line indicates the threshold of five MGEs/community.

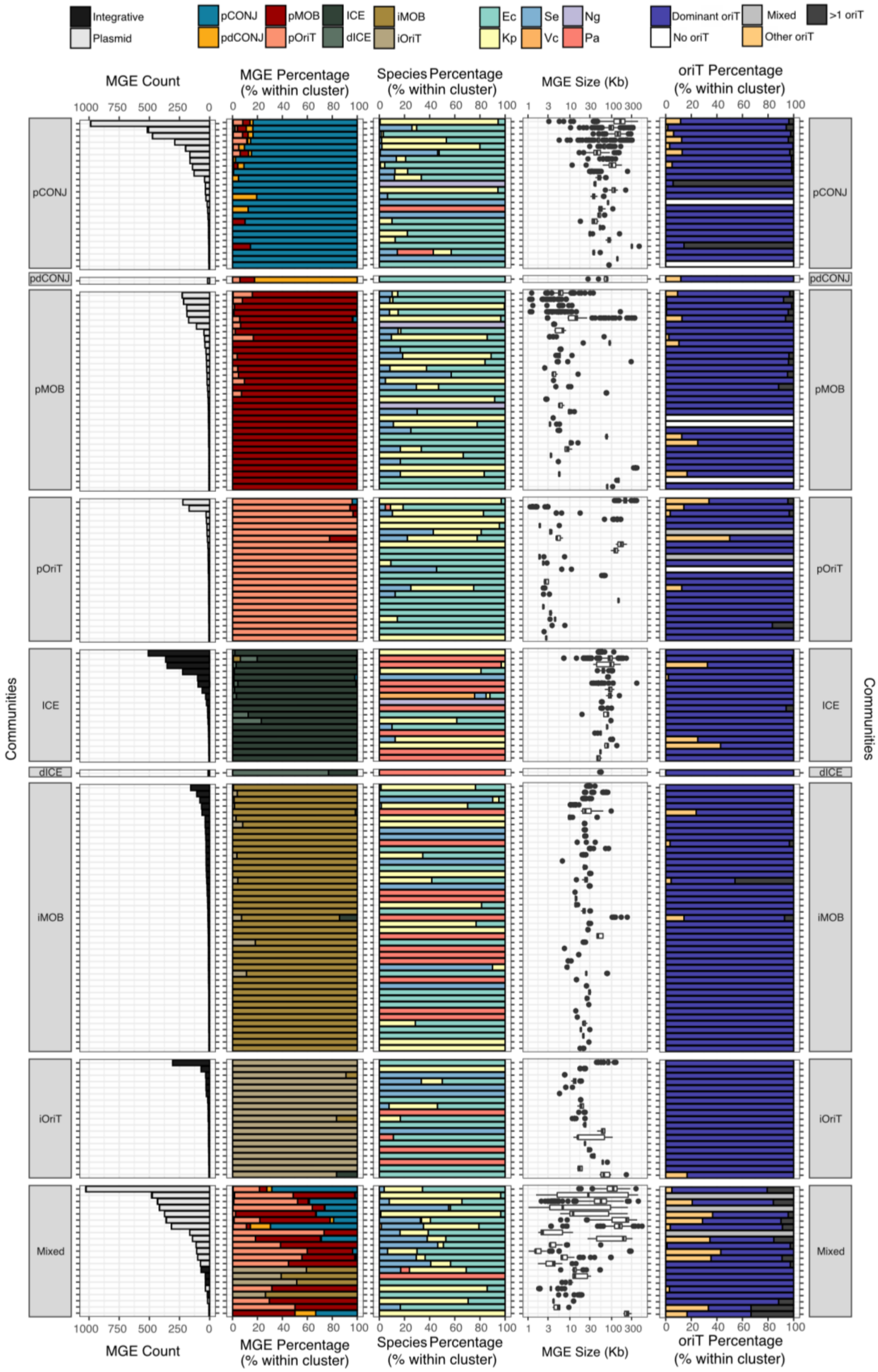
Integrative
Plasmid
pCONJ
pdCONJ
pMOB
pOriT
ICE
dICE
iMOB
iOriT
Ec
Kp
Se
Vc
Ng
Pa
Dominant oriT
No oriT
Mixed
Other oriT
>1 oriT
MGE Count
MGE Percentage (% within cluster)
Species Percentage (% within cluster)
MGE Size (Kb)
oriT Percentage (% within cluster)
Communities
pCONJ
pdCONJ
pMOB
pOriT
ICE
dICE
iMOB
iOriT
Mixed

104

**Figure S14.** Communities identified across the MGEs. The facets indicate the dominant MGE of the community (>75%). The first plot indicates the total number of MGEs in the community: light grey if plasmid, black if integrative element. The second plot indicates the distribution of each MGE type per community. The third plot indicates the distribution of bacteria hosts per community (species). The fourth plot indicates the size distribution of the MGEs of the community (log scale). The last plot indicates the *oriT* distribution. Blue when it is the dominant *oriT* (>75%). The legend of each plot is represented at the upper part of the figure.

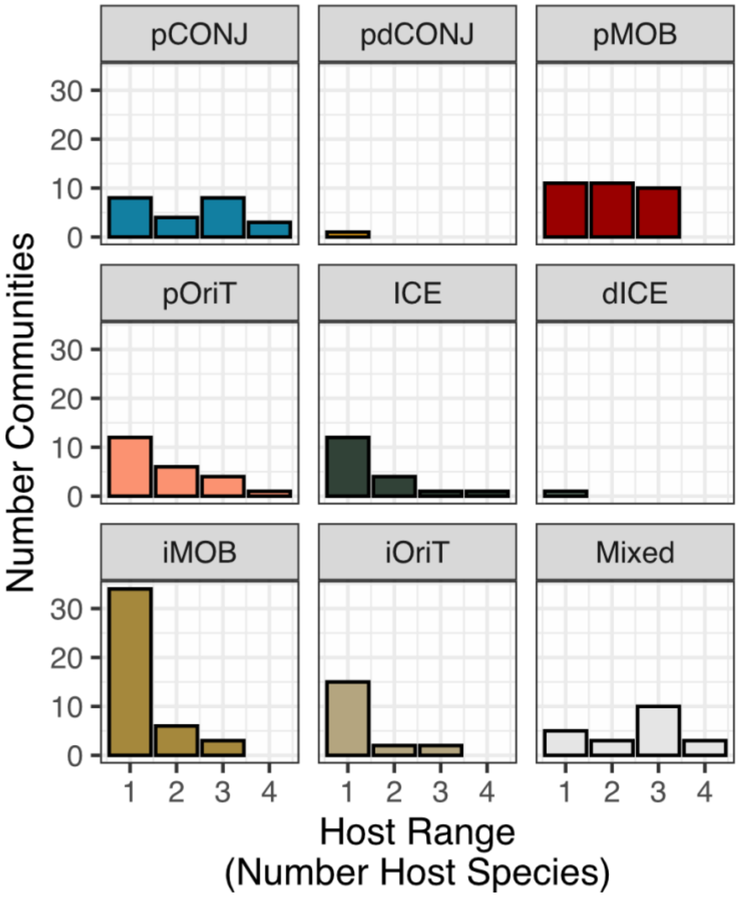


**Figure S15.** Host range of the MGE communities, measured as the number of different host species in which the community is present. Communities have been split into different facets according to its dominant MGE.

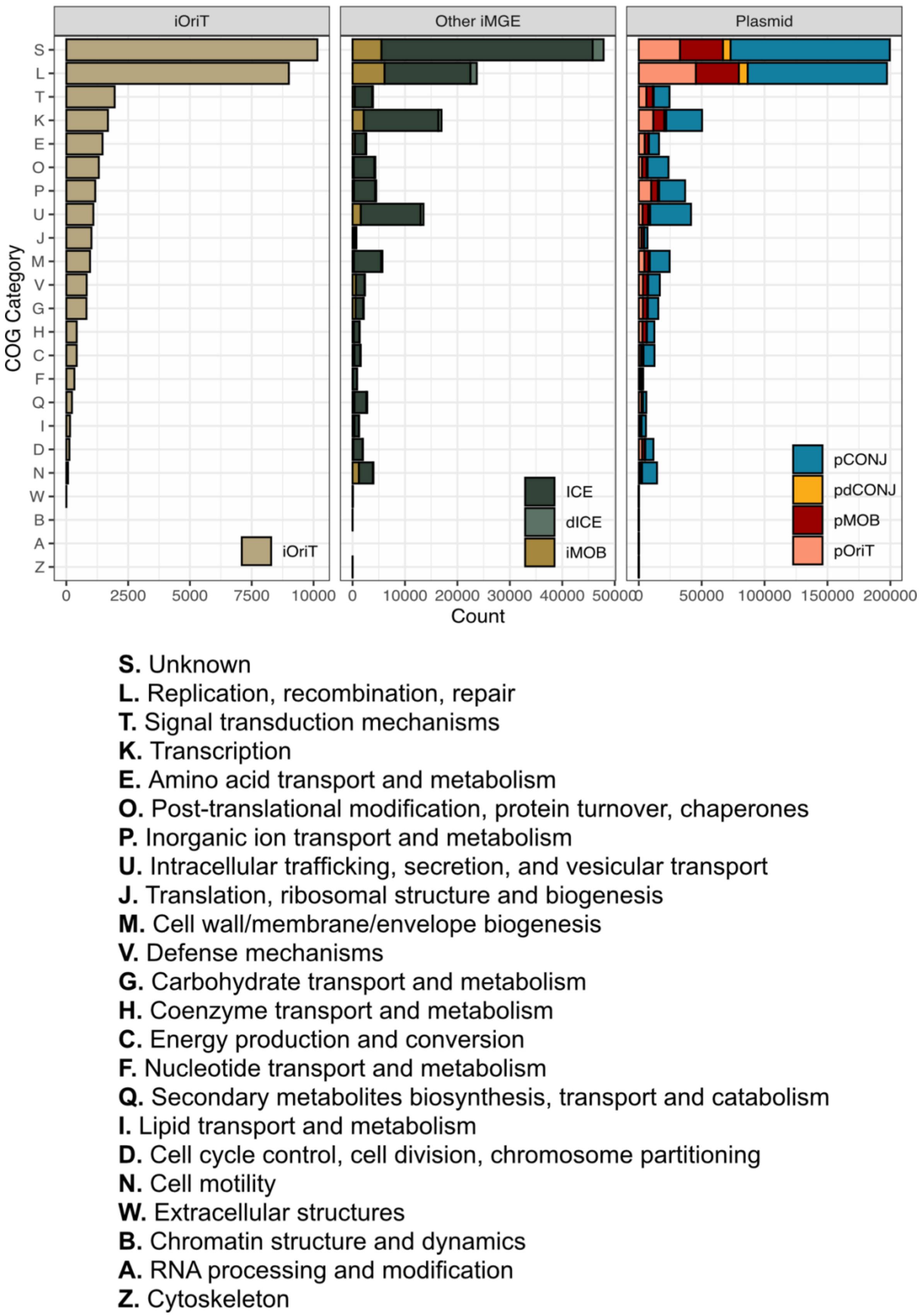


**Figure 16.** COG categories identified across the protein coding genes of the iOriTs (left), other integrative MGEs (middle), and plasmids (right). The colors indicate the MGE class.

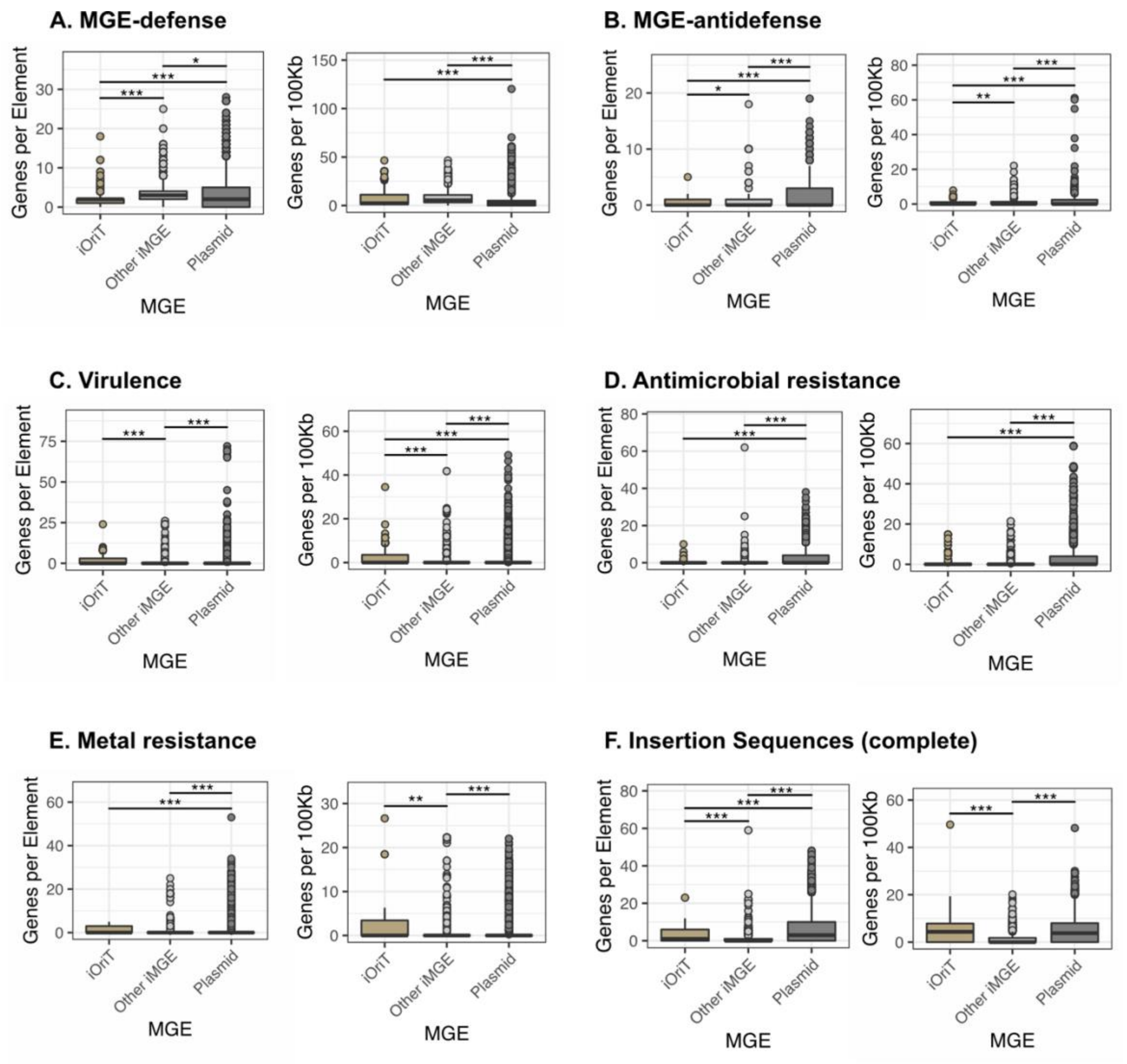


**Figure S17.** Number of genes per element (left) and genes per 100Kb (right) of functions associated to MGE-defense (**A.**), MGE-antidefense (**B.**), virulence (**C.**), antimicrobial resistance (**D.**), metal resistance (**E**.), and insertion sequences (**F.**). Asterisks indicate a statistically significant pairwise difference determined by Tukey's HSD post-hoc test: p-value < 0.001 (***), p-value < 0.01 (**), p-value < 0.05 (*).

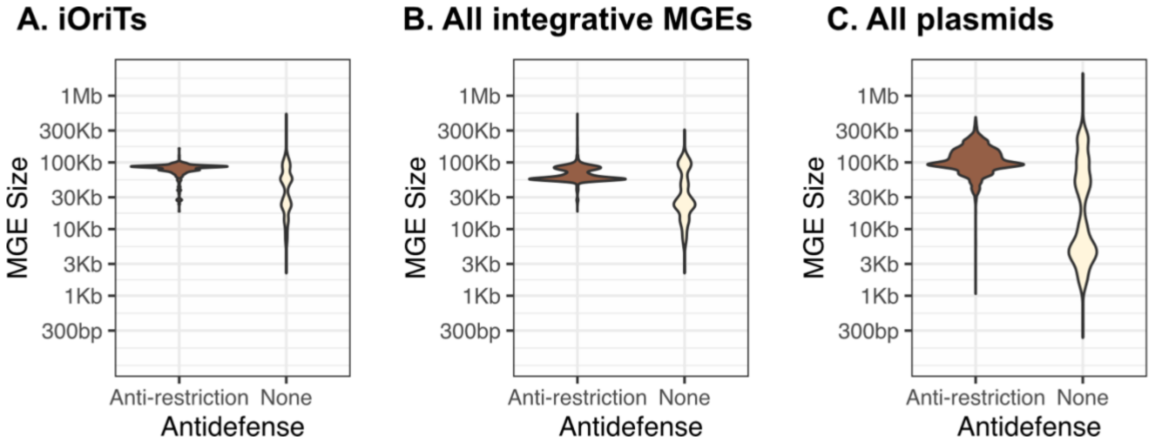


**Figure S18.** Size distribution of MGEs encoding or missing anti-restriction genes. **A.** iOriT elements. **B.** All integrative MGEs. **C.** Plasmids.

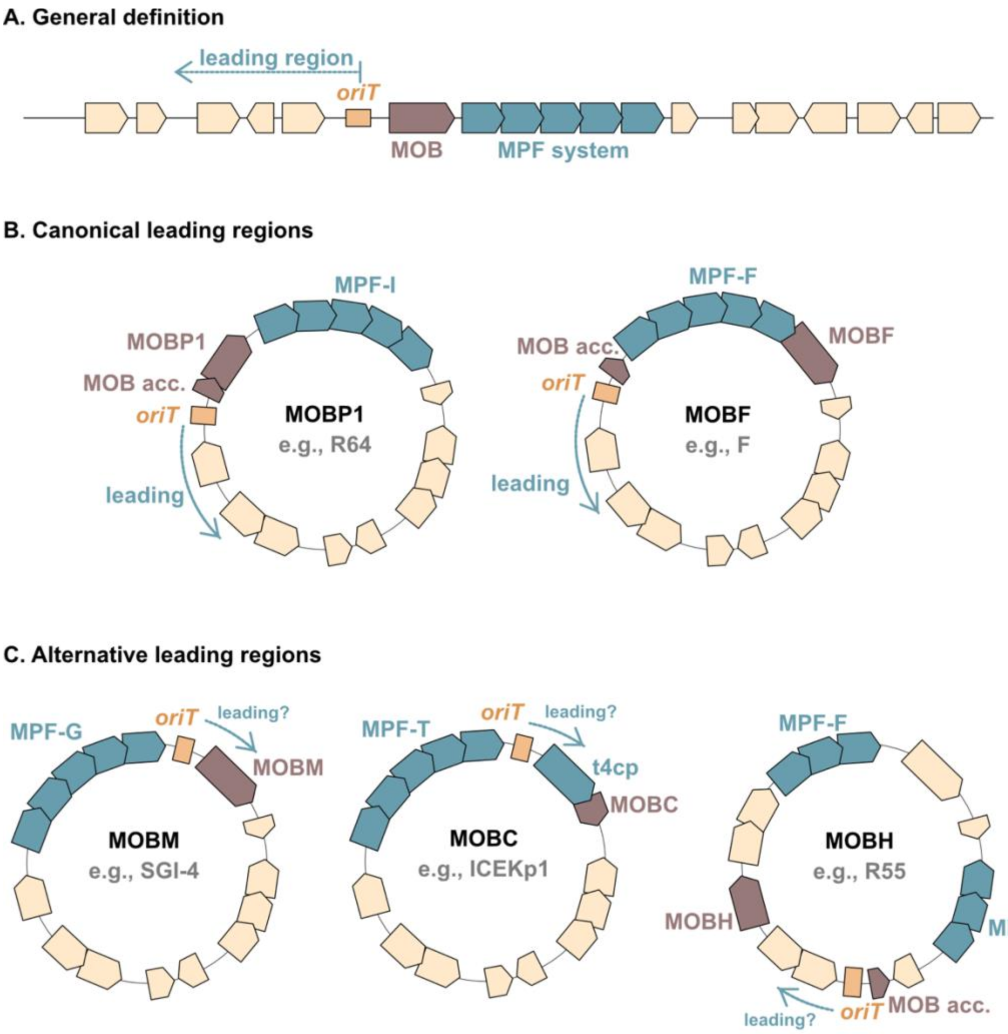


**Figure S19. A.** General definition of conjugative leading region. **B.** Conjugative machineries with leading regions matching the general definition. **C.** Alternative cases. MOB: relaxase. MPF: Mating Pair Formation system. MOB acc: relaxosome accessory. *oriT*: origin of transfer.

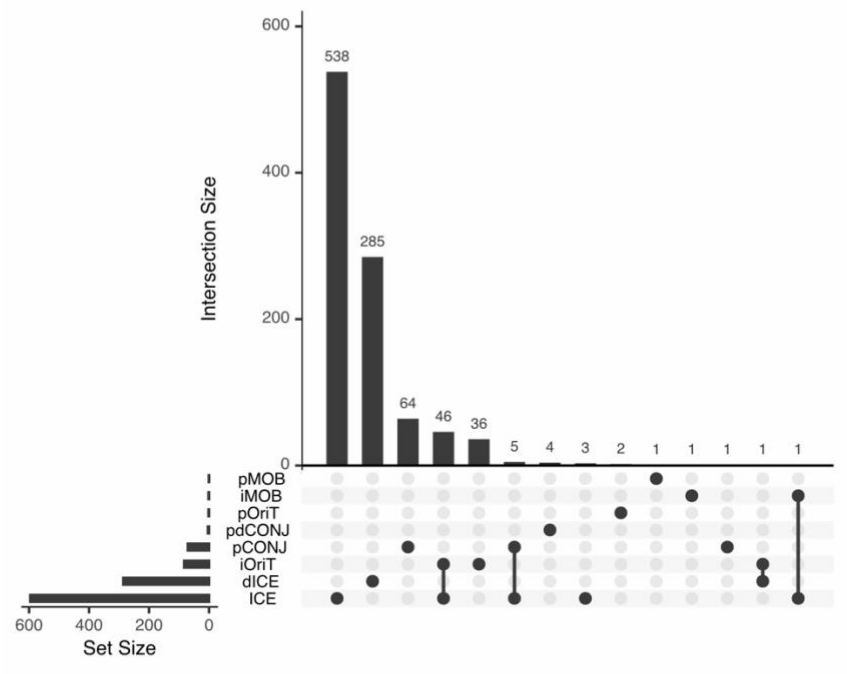


**Figure S20.** Co-occurrence of MGEs carrying an *oriT* of the family ICEK1 within the same genome. In case of integrative MGEs, we considered the interval-delimited Genomic Islands.

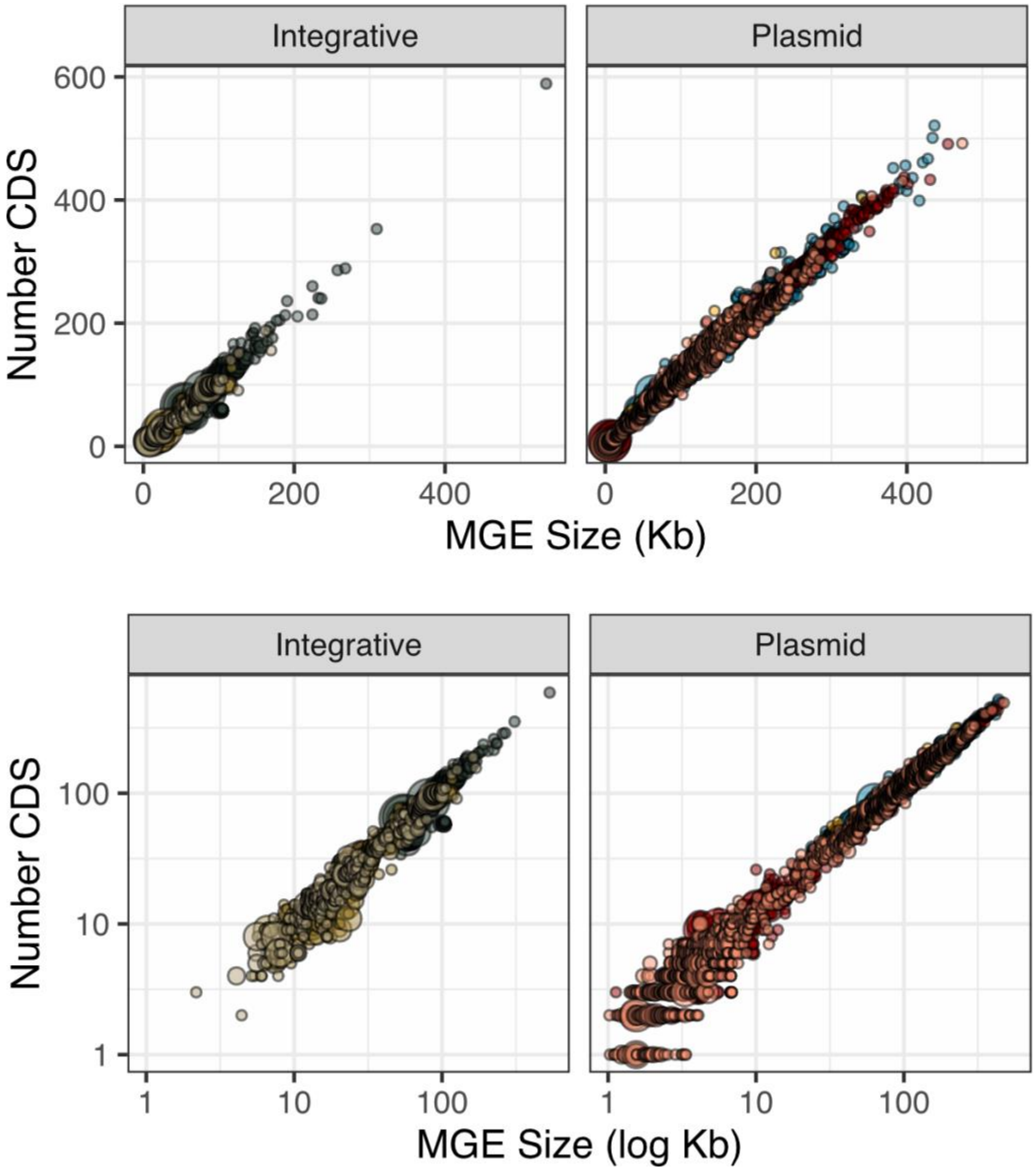


**Figure S21.** Number of protein coding sequences encoded by each MGE compared to its size (up: non-logarithmic scale; bottom: logarithmic scale), for both integrative MGEs (left) and plasmids (right). The color of each dot represents the MGE type, being the legend at the right of the figure. The width of the point is the number of elements with same size and gene number.
